# Cardiac Differentiation of Human Pluripotent Stem Cells Using Defined Extracellular Matrix Proteins Reveals Essential Role of Fibronectin

**DOI:** 10.1101/2021.04.09.439173

**Authors:** Jianhua Zhang, Ran Tao, Pratik A. Lalit, Juliana L. Carvalho, Yogananda Markandeya, Sean P. Palecek, Timothy J. Kamp

## Abstract

Research and therapeutic applications using human pluripotent stem cell-derived cardiomyocytes (hPSC-CMs) require robust differentiation strategies. Efforts to improve hPSC-CM differentiation have largely overlooked the role of extracellular matrix (ECM). The present study investigates the ability of defined ECM proteins to promote hPSC cardiac differentiation. Fibronectin, laminin-111, and laminin-521 enabled hPSCs to attach and expand; however, fibronectin ECM either endogenously produced or exogenously added promoted, while laminins inhibited, cardiac differentiation in response to growth factors Activin A, BMP4, and bFGF. Inducible shRNA knockdown of fibronectin prevented Brachyury^+^ mesoderm formation and subsequent hPSC-CM differentiation. Antibodies blocking fibronectin binding to integrin β1, but not α5, inhibited cardiac differentiation. Furthermore, inhibition of integrin-linked kinase blocked cardiac differentiation. These results identify fibronectin, laminin-111 and laminin-521 as defined substrates enabling cardiac differentiation of hPSCs and uncover the essential role of fibronectin and downstream signaling pathways in the early stage of hPSC-CM differentiation.

## Introduction

Human pluripotent stem cells (hPSCs) including embryonic stem cells (ESCs) and induced pluripotent stem cells (iPSCs) provide a powerful model system for research and therapeutic applications. Cardiomyocytes (CMs) derived from hPSCs have been widely and increasingly used in research for cardiac development and clinical applications for cardiac repair and regeneration, drug testing and precision medicine. Methods and protocols to differentiate hPSCs to CMs (hPSC-CMs) have been advanced significantly in the past decade. The most investigated cardiac differentiation protocols have focused on using soluble molecules including growth factors and small molecules to treat hPSCs to promote stage-specific cardiac progenitors and ultimately to differentiate to hPSC-CMs. These protocols also require extracellular matrix (ECM) as the substrate to enable hPSC attachment, survival, proliferation and differentiation. However, the ECM proteins involved in cardiac differentiation of hPSCs and the ECM activated signaling pathways have been far less investigated and elucidated. The ECM composition during development is dynamic and provides key signals contributing to stage-specific transitions [1]. Our previous study showed that hPSCs cultured on the commercially available ECM preparation, Matrigel, more efficiently and reproducibly differentiate to hPSC-CMs in response to Activin A/BMP4/bFGF signaling if they concurrently received overlays of Matrigel during the initiation of differentiation, the matrix sandwich protocol [2]. The Matrigel overlays promote the first stage of differentiation, the epithelial to mesenchymal transition to form Brachyury^+^ precardiac mesoderm, mimicking primitive streak in development [3]. However, Matrigel is a complex mixture of ECM proteins produced from Engelbreth-Holm-Swarm mouse sarcoma cells, is not fully defined, and exhibits batch-to-batch variability. The essential ECM components responsible for promoting the initial stages of cardiogenesis in the Matrigel sandwich protocol as well as the optimal ECM environment to promote cardiogenesis in general remain to be determined.

Complex mixtures of ECM proteins such as Matrigel have enabled attachment and self-renewal of hPSCs in appropriate media, and more recently recombinant ECM proteins and synthetic substrates have been identified that can support long-term culture of hPSCs [4]. These defined substrates mimic the ECM components present in the earliest embryo including laminins, collagens, fibronectin, vitronectin, and proteoglycans. The hPSCs interact with the substrates via the transmembrane receptors, integrins, and cell adhesion molecules, such as cadherins.

However, for cardiac differentiation protocols, a substrate that both allows attachment of the hPSCs and also supports proliferation and subsequent differentiation is needed. Strong signals to maintain self-renewal and pluripotency provided by the ECM will impede the differentiation processes, so a composition of ECM that is dynamic and plays multiple roles in support of hPSC proliferation as well as differentiation is theoretically optimal. Yap and colleagues utilized a combination of recombinant laminins, laminin-521 (LN521) which has been demonstrated to promote self-renewal of hPSCs [5] and laminin-221 (LN221) to enable differentiation to cardiac progenitors [6]. Others using a design of experiment statistical approach found a combination of three ECM proteins optimal for cardiac differentiation of hPSCs, collagen type I, laminin-111 (LN111) and fibronectin (FN) [7, 8]. Burridge and colleagues systematically tested a range of different substrates in a defined small molecule-based cardiac differentiation protocol and found a variety of substrates including a synthetic vitronectin-derived peptide, recombinant E-cadherin, recombinant human vitronectin, recombinant human LN521, truncated LN511 and human FN, and a FN mimetic enabled hPSC-CM differentiation [9]. However, these important studies have not examined the impact of dynamic manipulation of defined ECM proteins or ECM signaling pathways in cardiac differentiation, nor characterized the changes in endogenous ECM proteins that ultimately contribute to the cellular transitions. In the present study, we tested a variety of human recombinant and defined ECM proteins for both attachment and overlay of hPSC cultures in the matrix sandwich protocol and investigated the ECM activated signaling pathways. We found that of the tested ECM proteins, only FN overlays promoted cardiac differentiation comparable to Matrigel overlays; in contrast, LN111, LN521 and collagen IV (COL4) overlays inhibited cardiogenesis. Furthermore, hPSCs differentiated efficiently to hPSC-CMs without overlay when grown on FN, LN111, and LN521. Regardless of the ECM preparation used as the attachment substrate, we identified an essential role of FN promoting the initial stages of differentiation acting via transmembrane integrin β1 receptors and potential downstream integrin-linked kinase (ILK) signaling.

## Results

### Defined extracellular matrix proteins support hPSC adhesion, growth and cardiac differentiation

We tested whether defined human ECM proteins could replace Matrigel in the matrix sandwich cardiac differentiation protocol. The matrix sandwich protocol uses Matrigel coating for hPSC adhesion and expansion followed by overlaying the proliferating hPSCs with Matrigel at day -2 and day 0 of differentiation with the addition of growth factors Activin A at day 0-1, followed by addition of BMP4 and bFGF at day 1-5 (Fig. 1A). Therefore, we first tested the ability of defined ECM proteins to support hPSC adhesion and expansion. DF19-19-11T iPSCs or H1 ESCs were seeded on human LN111, LN521, COL4, and FN coated surfaces and cultured in mTeSR1 medium. The hPSCs grew as a monolayer on LN111, LN521, and FN and exhibited similar morphology and expression of pluripotency markers of OCT4 and SSEA4 as hPSCs grown on Matrigel (Fig. 1B and Supplementary Fig. S1). However, the hPSCs seeded on COL4 did not grow as a confluent monolayer (Supplementary Fig. S2), so this matrix coating was not tested further.

**Fig. 1.**
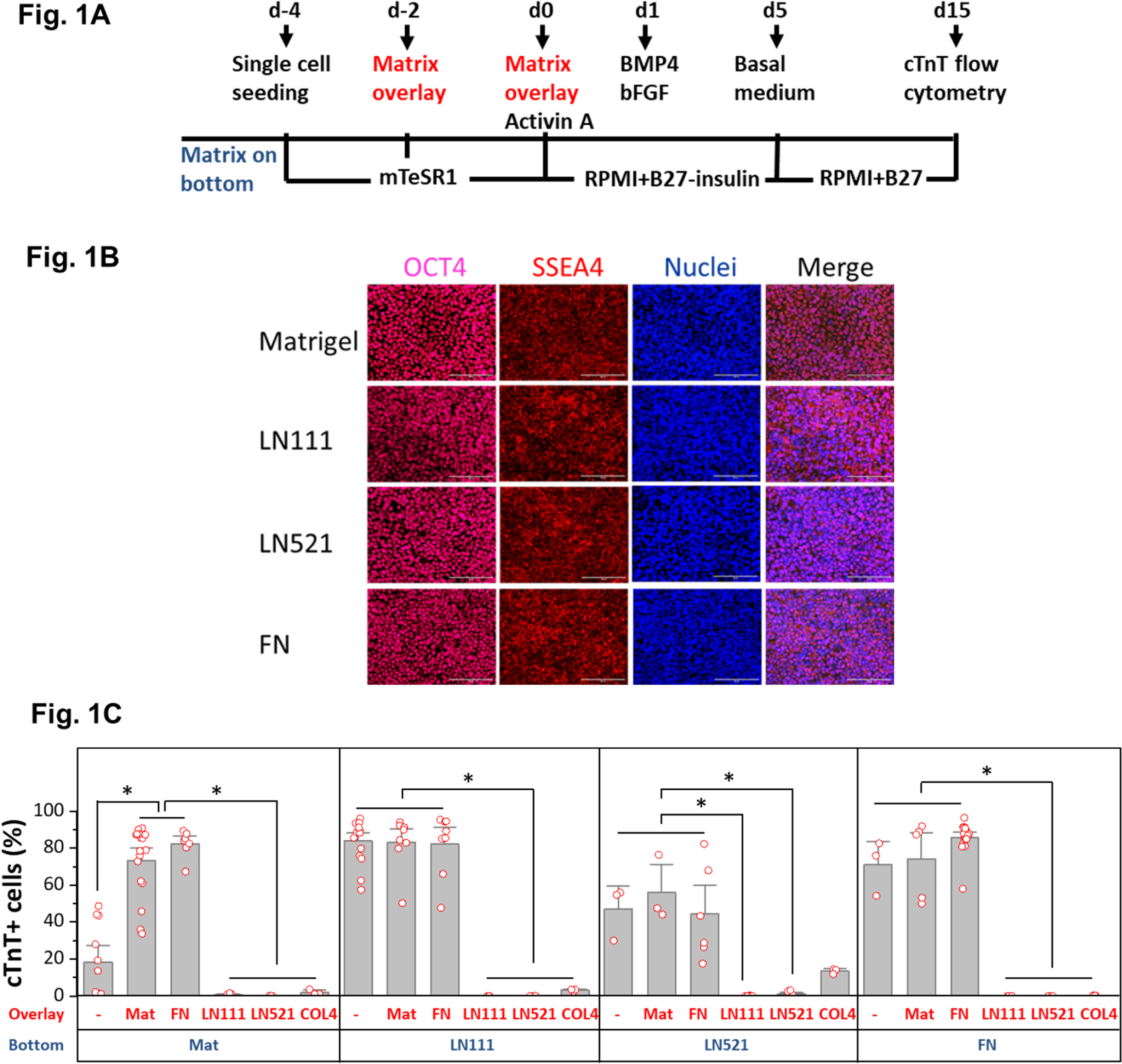
Defined ECM proteins support hPSCs adhesion, growth and cardiac differentiation using the matrix sandwich protocol. (A) Schematic method of the matrix sandwich protocol. Defined ECM proteins are tested as coating matrix and overlay matrix in RED. (B) Fluorescence images of DF19-9-11T iPSCs growing on the ECM of Matrigel, LN111, LN521 and FN as confluent monolayer and immuno-labeled with antibodies against OCT4 and SSEA4. Scale bar is 100 µm. (C) cTnT^+^ cells measured by flow cytometry at 15 days of differentiation of DF19-9- 11T iPSCs on different ECM proteins as substrate (bottom) and overlay (overlay). N ≥ 3 biological replicates. Error bars represent SEM. *P<0.05, one-way ANOVA with post-hoc Bonferroni and Tukey test.

For those matrix substrates which supported monolayer growth of hPSCs, including LN111, LN521 and FN, we tested overlay of defined ECM proteins in the matrix sandwich protocol. Cardiac differentiation was measured by flow cytometry for cTnT^+^ cells at 15 days of differentiation (Fig. 1C and Supplementary Fig. S3). DF19-9-11T iPSCs seeded on Matrigel coated surface showed poor cardiac differentiation in response to the growth factors without an overlay of matrix, but with Matrigel overlays the percentage of cTnT^+^ cells was significantly increased as we previously reported [2]. Interestingly, FN overlays were as effective as Matrigel overlays in promoting cardiac differentiation of the hPSCs growing on Matrigel. However, if cells were seeded on LN111, LN521, or FN coated surfaces, the overlay of Matrigel or FN did not further increase the efficiency of hPSC-CM generation, and overlays of LN111, LN521 and COL4 strongly inhibited cardiac differentiation. These results demonstrated that the defined ECM proteins of LN111, FN and to a lesser extent LN521 support hPSC adhesion, growth and cardiac differentiation in monolayer hPSC culture and do not require a matrix overlay for efficient cardiac differentiation using the Activin A/BMP4/bFGF growth factor-directed protocol.

### hPSCs monolayer culture on LN111 substrate promoted endogenous FN production

Because FN promoted cardiac differentiation as coated matrix protein as well as an overlay matrix protein and because FN plays key roles in EMT during gastrulation and cardiogenesis [3, 10–21], we next examined for the presence of FN ECM in hPSC culture. DF19-9-11T iPSCs plated on the Matrigel-coated surface and cultured in mTeSR1 per the matrix sandwich protocol were immunolabeled with a FN antibody on day -3, -2, -1 and 0 without permeabilizing the cells to examine extracellular FN protein (Fig. 2A). On day -3 and -2, minimal immunolabeled FN in ECM was observed. However, after 4 days of culture (day 0) immunolabeled fibrillar FN ECM was abundant in the matrix sandwich culture (Fig. 2A, B). In contrast, the monolayer culture control in the absence of Matrigel overlay had significantly less FN ECM present by day 0 (Fig. 2A, B). This suggests that the Matrigel overlay promotes production of endogenous FN or remodeling of FN ECM relative to the monolayer culture control without Matrigel overlay. Because hPSC cultured on LN111 coated surface without matrix overlay enabled efficient cardiac differentiation (Fig. 1C), we examined the endogenous FN production in the hPSC culture on LN111 coated surface in which no exogenous FN was added. DF19-9-11T iPSCs growing on LN111 coated surface without any matrix overly were immunolabeled with the FN antibody without permeabilizing the cells. Confocal z-scan of the cell culture showed no detectable FN ECM at day -3 and -2, similar to the Matrigel/Matrigel sandwich culture; however, by day 0, dense fibrillary FN ECM was present in the cell culture on LN111 coated surface (Fig. 2C). Similar to our previous study of the Matrigel/Matrigel sandwich culture [2], the hPSCs growing on the LN111 coated surface without any matrix overlay formed multilayer cultures as shown in the side view of the confocal z-scan as did FN/FN matrix sandwich culture (Fig. 2D). To determine if the hPSC culture results in accumulation of endogenously produced laminin ECM as well, DF19-9-11T iPSCs cultured on FN, LN111 and LN521 coated surfaces were immunolabeled with an antibody detecting laminins without permeabilizing the cells, and did not show measurable laminin ECM after 4 days of growth on these defined matrices (Supplementary Fig. S4). These results together with the above cardiac differentiation results supported the potential role of FN in promoting ActivinA/BMP4/bFGF directed hPSC cardiac differentiation.

**Fig. 2.**
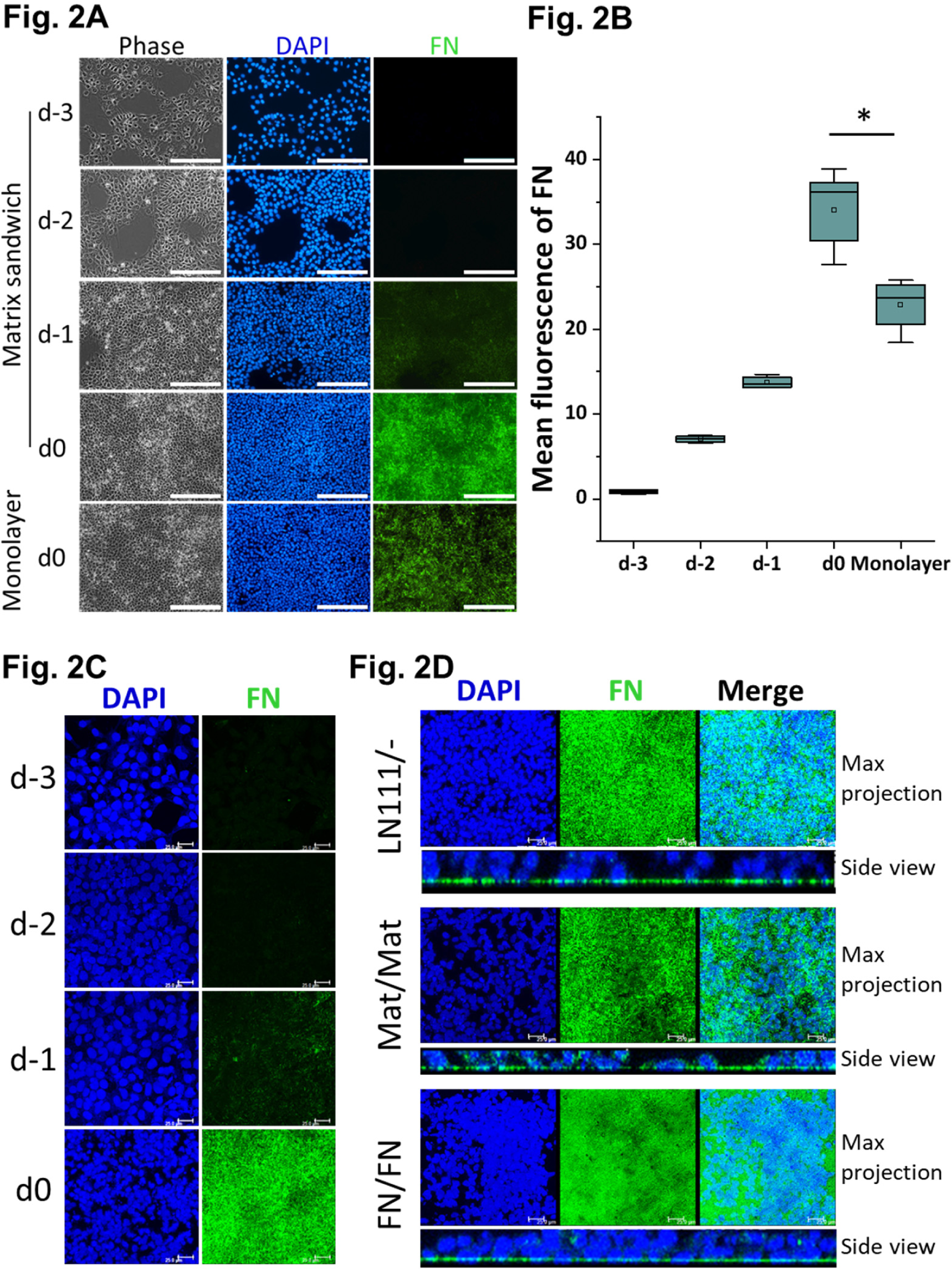
Production of endogenous FN in the hPSC-matrix sandwich and LN111 culture. (A) Phase contrast and fluorescence images of the matrix sandwich culture of DF19-9- 11T iPSCs growing for 4 days and immunolabeled with fibronectin antibody, compared with the monolayer culture. Scale bar is 200 µm. (B) Quantitative analysis of the FN fluorescence in A by Image J. N ≥ 3 replicates. The box plots summarize the biological replicates with the box enclosing from first to third quartile and middle square indicating mean and line in box indicating median. *P<0.05, one-way ANOVA with post-hoc Bonferroni and Tukey test. (C) Maximum projection view of the confocal z-scan of DF19-9-11T iPSCs growing for 4 days on LN111 coated surface immunolabeled with fibronectin antibody. Scale bar is 25 µm. (D) The maximum projection view and side view of the confocal z-scan of DF19-9-11T iPSCs growing for 4 days on LN111 coated surface without matrix overlay and immunolabeled with the fibronectin antibody. The multiplayer culture and FN production are similar as in Matrigel/Matrigel and FN/FN matrix sandwich culture. Scale bar is 25 µm.

### Differentiation of hPSCs on LN111 substrate undergo EMT and generate precardiac mesoderm in FN rich ECM

To characterize the early stages of cardiac differentiation of hPSCs cultured on LN111 and treated with Activin A/BMP4/bFGF signaling, we examined markers of EMT, mesoderm and cardiac mesoderm. Gene expression was assessed by quantitative RT-PCR, and upon the addition of Activin A, BMP4 and bFGF, there was significant upregulation of transcription factors associated with EMT including *SNAI1* [22], *SNAI2* [23] and *TWIST* [24] (Fig. 3A). The mesenchymal cell markers of vimentin (*VIM*), fibronectin (*FN1*) and N-cadherin (*CDH2*) were also greatly upregulated by day 3 (Fig. 3A). In contrast, E-cadherin expression (*CDH1*), an epithelial cadherin, was greatly downregulated by day 3 of differentiation. The mesendoderm/precardiac mesoderm transcription factors *GSC*, *MIXL1*, *SOX17*, *TBXT* and *MESP1* were transiently upregulated followed by expression of cardiac transcription factors of *ISL1*, *NKX2-5* and *GATA4* at day 3-5 (Fig. 3B).

**Fig. 3.**
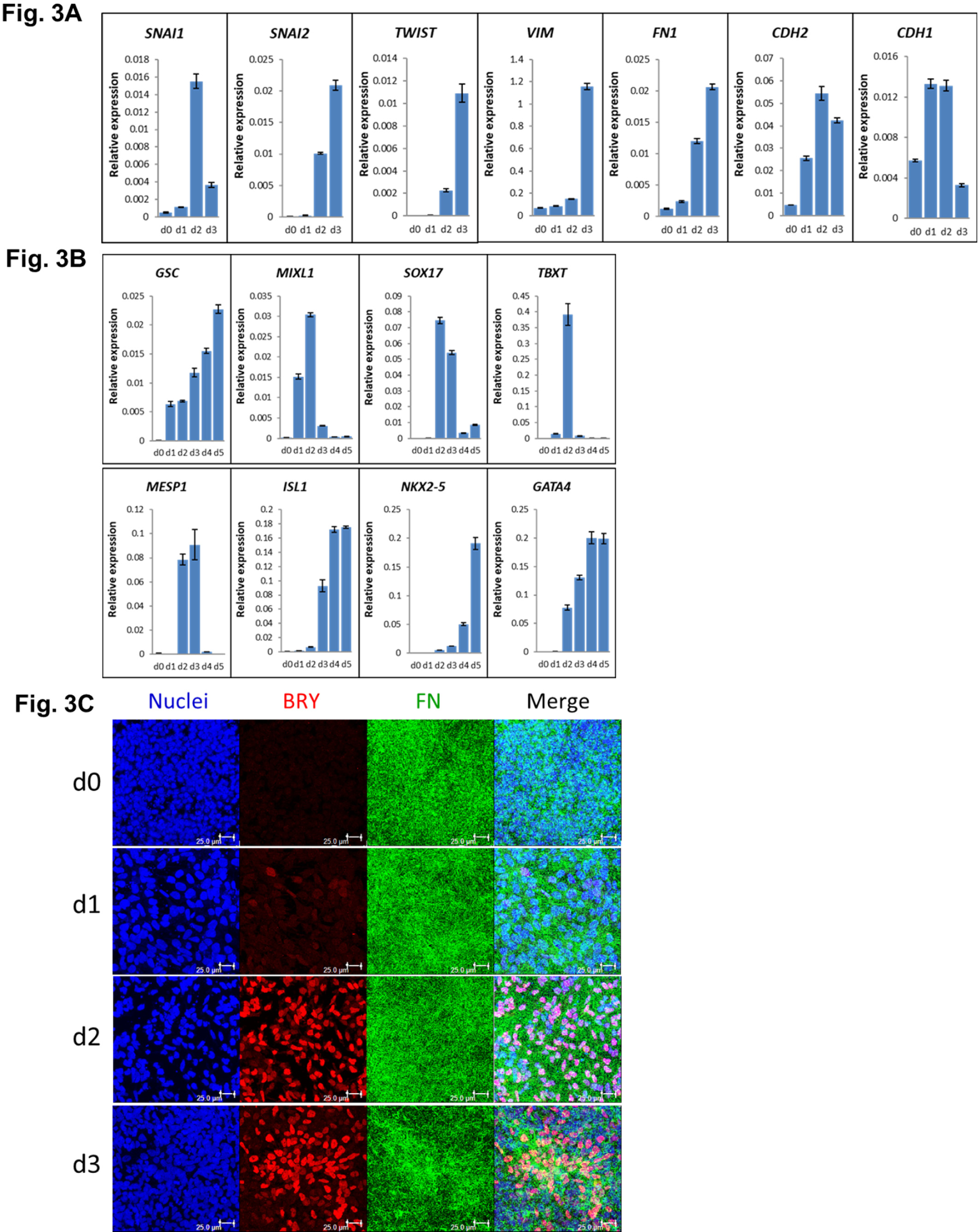
Expression of EMT and precardiac mesoderm markers in the cardiac differentiation of hPSCs cultured on LN111 substrate by Activin A/BMP4/bFGF signaling. (A) qRT-PCR for gene expression of EMT markers at day 0-3 of cardiac differentiation. (B) qRT-PCR for gene expression of precardiac mesoderm markers at day 0-5 of cardiac differentiation. N=3 technical replicates for each point. (C) Maximum projection view of the confocal z-scan of DF19-9- 11T iPSCs at day 0-3 of the cardiac differentiation co-labeled with antibodies against Brachyury (BRY) and Fibronectin (FN). Scale bar is 25 µm. Error bars represent SEM.

To determine if FN ECM persisted or remodeled during the early stages of cardiac differentiation on LN111 substrate, immunolabeling of the early differentiated DF19-9-11T iPSCs on day 0-3 for FN and Brachyury was performed. Confocal z-scan imaging showed abundant fibronectin ECM at each day, and Brachyury^+^ cells were associated with the dense network of FN ECM (Fig. 3C), suggesting that the Brachyury^+^ cells interact with FN. Similar Brachyury^+^ cells and the FN ECM network were also observed in the Matrigel/Matrigel and FN/FN matrix sandwich cultures (Supplementary Fig. S5). Together, these results show that hPSCs grown on LN111, undergo the early stages of cardiac differentiation with transitions to mesoderm and cardiac mesoderm progenitors occurring in an endogenously generated FN rich ECM, similarly as in the Matrigel/Matrigel and FN/FN matrix sandwich cultures.

### FN is essential for cardiac differentiation of hPSCs

To determine if FN is essential for precardiac mesoderm formation in our protocol and to investigate the stage-specific roles of FN during cardiac differentiation of hPSCs, we generated a doxycycline inducible FN knockdown system using *FN1* shRNA (Supplementary Fig. S6). The two vectors shown in Fig. 4A were incorporated into lentivirus and transduced into hPSCs. Clones were selected by neomycin resistance from both hESC line H1 and hiPSC line DF19-9- 11T. To confirm the doxycycline inducibility of the *FN1* shRNA in the cell lines, we first examined for doxycycline-induced bicistronic mCherry (Supplementary Fig. S7A). Inducible FN knockdown was demonstrated by immunolabeling with FN antibody (Supplementary Fig. S7B) and quantitative western-blot for FN expression (Supplementary Fig. S7C, D).

**Fig. 4.**
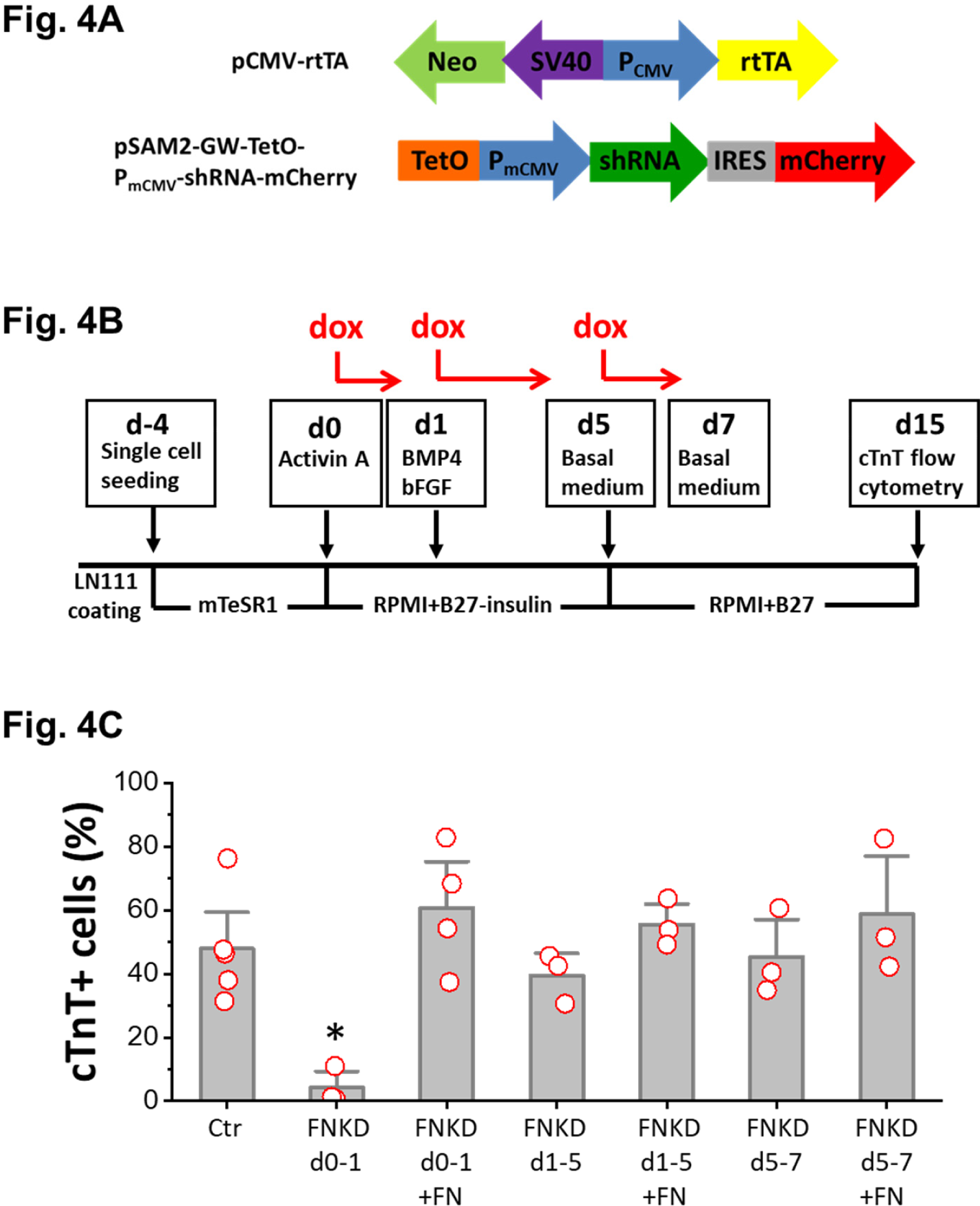
FN is essential for precardiac mesoderm formation in the cardiac differentiation of hPSCs. (A) Schematic of the inducible shRNA construct for *FN1* knockdown. (B) Schematic method of FN knockdown at differentiation stages of day 0-1, 1-5 and day 5-7 in the cardiac differentiation protocol. (C) cTnT^+^ cells measured by flow cytometry at 15 days of differentiation of the H1 *FN1* knockdown clone using the protocol in Fig. 4B. N ≥ 3 biological replicates. Error bars represent SEM. *P<0.05, one-way ANOVA with post-hoc Bonferroni and Tukey test. FNKD indicates FN knockdown by dox induction. +FN indicates exogenous FN added.

Cardiac differentiation was performed using the monolayer based protocol with hPSCs growing on LN111 coated surface and treated with growth factors of Activin A, BMP4 and bFGF as shown in Fig. 4(B) To probe the stage-specific effect of FN knockdown during cardiac differentiation, dox was added at different time points: day 0-1, day 1-5, and day 5-7, and cardiac differentiation was measured by flow cytometry for cTnT^+^ cells at 15 days of differentiation. Cardiac differentiation was significantly inhibited when FN was knocked down at day 0-1, whereas FN knockdown at day 1-5, or day 5-7, did not have significant impact on the cTnT^+^ cells when compared to the no dox control (Fig. 4C).

Because FN knockdown at day 0-1 significantly inhibited cardiac differentiation, we next tested if exogenous FN at this stage can rescue cardiac differentiation. Using the same protocol as shown in Fig. 4B with dox induction of FN knockdown at day 0-1, we added soluble human FN (3ug/cm^2^) in the cell culture on day 0-1. Cells were differentiated for 15 days, and cardiac differentiation was measured by flow cytometry for cTnT^+^ cells as above. Adding exogenous FN fully rescued the cardiomyocyte differentiation, giving rise to similar percentage of cTnT^+^ cells compared to the no FN knockdown control (Fig. 4C). However, adding exogenous FN (3ug/cm^2^) at day 1-5 or day 5-7 did not significantly increase the percentage of cTnT^+^ cells when compared to the no dox control at day 1-5 and day 5-7, respectively (Fig. 4C). Furthermore, the dox induction of FN knockdown is concentration dependent (Supplementary Fig. S8). With the dox concentration higher than 4ug/ml, it caused significant cell death after 24 hours and no cells survived by 15 days of differentiation, and the cardiac differentiation could not be rescued with adding exogenous FN, indicating the toxicity of doxcycline to the cells (Supplementary Fig. S8A). We confirmed the detrimental effect of the high concentration of dox when added at day 0-1 during the cardiac differentiation with the regular hPSC line in which high concentration of dox caused significant cell death and no cells survived after 2 days of differentiation (Supplementary Fig. S8B).

### FN is required for formation of Brachyury^+^ cells

Because FN knockdown at day 0-1 dramatically inhibited cardiac differentiation which could be rescued by exogenous FN, we first evaluated the expression of EMT genes which mark the initial transition of hPSCs to mesoderm over the first two days of differentiation. Quantitative RT-PCR was performed at 0, 24, 36 and 48h after hPSCs differentiation was initiated (Fig. 5A). The effect of dox-induced knockdown of *FN1* transcripts was confirmed by the greater than 50% reduction in mRNA levels in the FN knockdown and FN knockdown + exogenous FN cell samples at 24h relative to the no dox control (Fig. 5A). By 36h, *FN1* transcripts recovered to the control level for FN knockdown condition or were significantly increased in the FN knockdown + exogenous FN samples after dox was removed at 24h. By 48h, there were similar level of *FN1* transcripts in both the control and the FN rescue samples, but no cells survived in the FN knockdown group (Fig. 5A). The key EMT transcription factors upregulated during gastrulation [3, 20, 25, 26], *SNAI1 and SNAI2*, were examined. Quantitative RT-PCR showed *SNAI1* expression increased significantly in both the FN knockdown and FN knockdown + exogenous FN samples compared to the control at 24h, and its expression continuously upregulated in the FN knockdown sample by 36h. By 48h there were similar level of *SNAI1* expression in both the control and the FN rescue samples; whereas, *SNAI2* expression was not significantly different between the groups but gradually upregulated in all three groups in the time course (Fig. 5A). *VIM* expression, similar to *SNAI1* expression, was increased significantly in the FN knockdown cells by 36h compared to the control (Fig. 5A), which is consistent with this mesenchymal marker and known target of *SNAI1*.

**Fig. 5.**
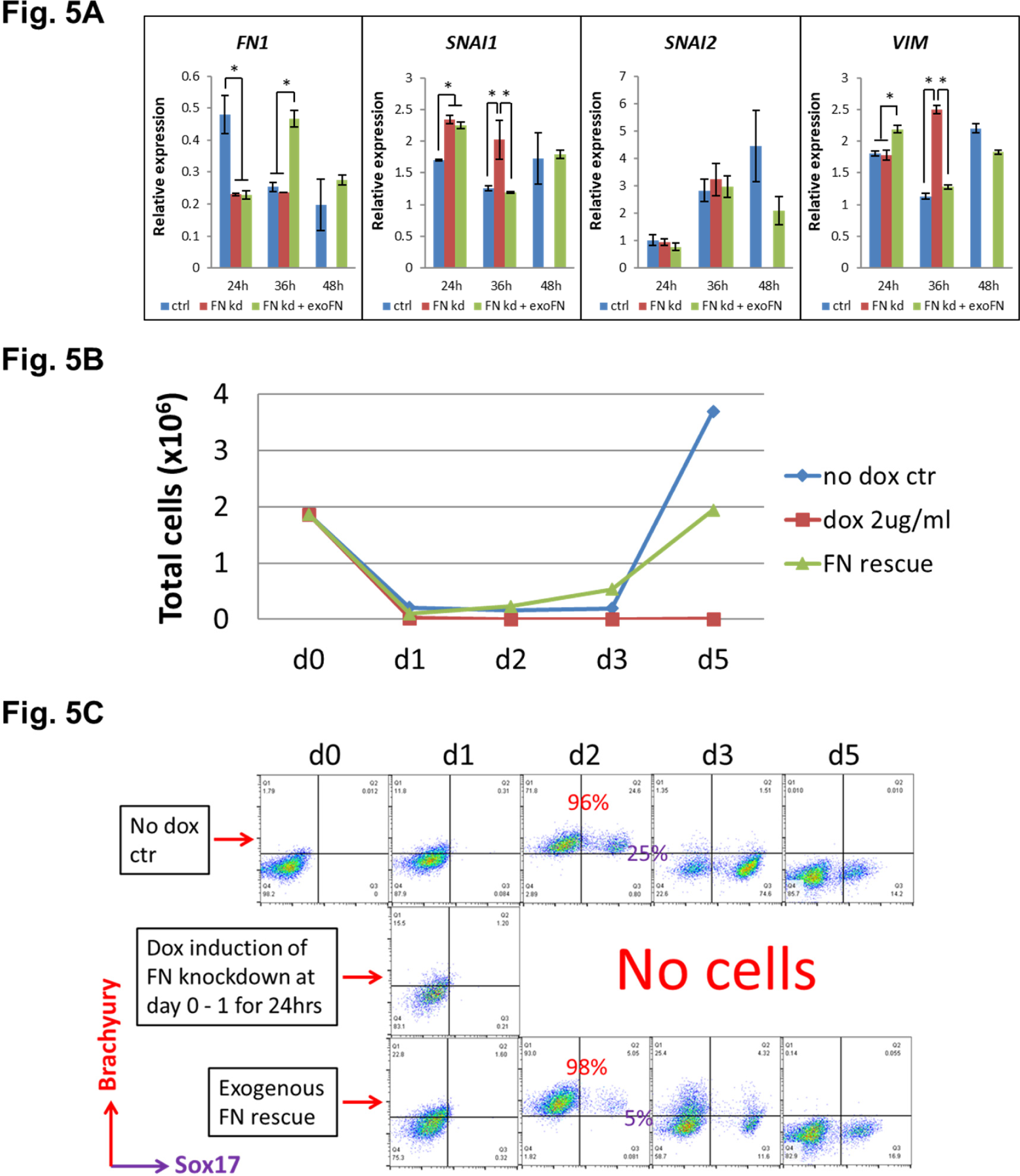
Knockdown FN at the precardiac mesoderm stage in the Activin A/BMP4/bFGF directed cardiac differentiation results in loss of Brachyury^+^ cells, exogenous FN can rescue it. (A) qRT-PCR for gene expression of EMT markers for the H1 FN knockdown clone in the cardiac differentiation time course of 0-48h at the no dox control, dox induction at day 0-1 and dox induction at day 0-1 with adding exogenous FN conditons. (B) Total cell number of the H1 FN knockdown clone in the time course of day 0-5 in the cardiac differentiation at the no dox control, dox induction at day 0-1 and dox induction at day 0-1 with adding exogenous FN conditions. (C) Flow cytometry of co-labeling the cells shown on with Brachyury and Sox17 antibodies. Error bars represent SEM. *P<0.05, one-way ANOVA with post-hoc Bonferroni and Tukey test.

We next examined the mesendodermal/precardaic mesodermal progenitors generated in the initial differentiation stage by flow cytometry. The cell counts of the attached cells on day 0-5 of differentiation showed a great reduction of cell number at day 1 in all three groups (Fig. 5B); however, the cells in the control and the FN rescue groups rapidly proliferated after day 1. In contrast, no cells survived in the FN knockdown group after day 2 (Fig. 5B). Because Brachyury and Sox17 are both expressed in mesendodermal progenitors, we co-labeled the cells with Brachyury and Sox17 antibodies on day 0-5 and analyzed by flow cytometry. Brachyury^+^ cells started to emerge at day 1 in all three groups. By day 2, 96-98% of the cells were Brachyury^+^ in both the control and FN rescue groups, but there were no surviving cells in the FN knockdown samples (Fig. 5C). The fraction of cells that were Brachyury^+^ rapidly decreased after day 2 in both the control and FN rescue groups, and by day 5 there were minimal Brachyury^+^ or Sox17^+^ cells present in both groups (Fig. 5C). We performed the same experiment using the DF19-9-11T FN knockdown clones and observed similar results (Supplementary Fig S9). In contrast to the day 0-1 treatment with dox, a later timed dox pulse from day 1-5 did not alter the abundance of Brachyury^+^ cells in all three groups over the same time course of differentiation (Supplementary Fig. S10). Together these results suggest that knockdown of FN at day 0-1 does not stop the initiation of EMT upon addition of Activin A at d0 [3, 20], but it prevents the generation and/or survival of Brachyury^+^ precardiac mesodermal progenitors. In contrast, shRNA knockdown of FN at later time points in the protocol did not impact the fate of the differentiating cells.

### FN acts via integrin β1 and integrin-linked kinase signaling to promote cardiac differentiation

To investigate the downstream signaling pathways contributing to FN’s essential role in formation of precardiac mesoderm, we investigated integrin binding and integrin-linked kinase (ILK) signaling. FN preferentially interacts with α5β1, αvβ1 and αvβ3 integrin heterodimers on the surface of cells [27, 28]. We first tested blocking integrin α5 with a monoclonal antibody, P1D6, or integrin β1 with a monoclonal antibody, P5D2, at day -2 or day 0 in the matrix sandwich protocol (Fig. 6A). Adding the antibody P1D6 that binds integrin α5 at day -2 or day 0 when Matrigel overlays were applied did not block cardiac differentiation as measured by flow cytometry of the cTnT^+^ cells (Fig. 6B). However, adding antibody P5D2 that binds integrin β1 at day 0 significantly blocked cardiac differentiation (Fig. 6B). Furthermore, the P5D2 antibody resulted in concentration-dependent inhibition of cardiomyocyte differentiation with the highest concentration tested (5 ug/ml) completely blocking cardiomyocyte differentiation (Fig. 6C).

**Fig. 6.**
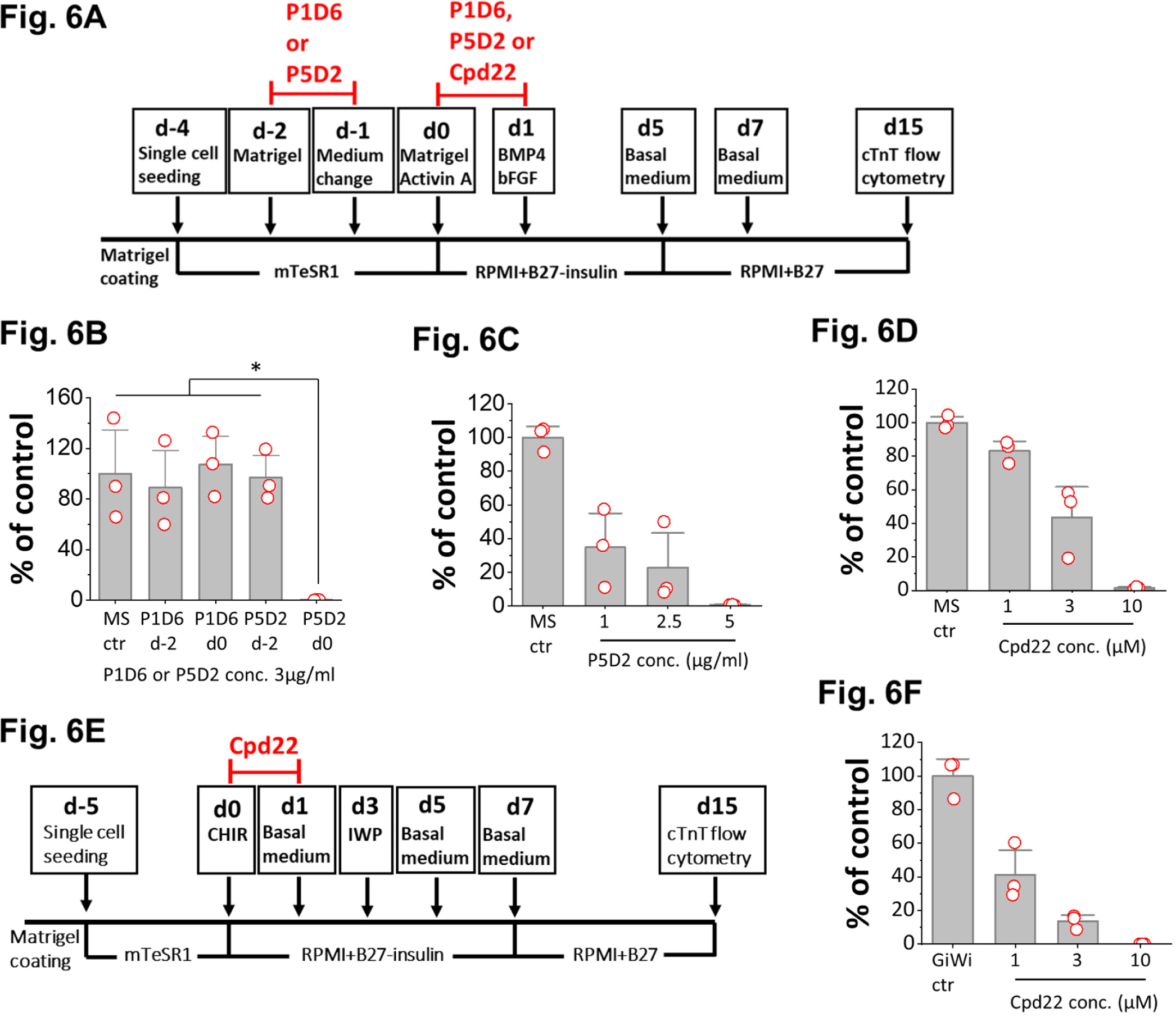
Cardiac differentiation is blocked by integrin β1 antibody and ILK inhibitor when added at day 0 in the matrix sandwich and GiWi protocols. (A) Schematic for testing monoclonal antibodies to block integrins α5 and β1 (P1D6 and P5D2, respectively) and inhibition of ILK by small molecule inhibitor Cpd22 in the matrix sandwich (MS) protocol. (B) cTnT^+^ cells measured by flow cytometry at 15 days differentiation when anti-human integrin α5 (P1D6) or β1 (P5D2) antibody was added at day -2 or day 0 at 3 µg/ml in the matrix sandwich protocol as shown in A. (C) cTnT^+^ cells measured by flow cytometry at 15 days differentiation when anti-human integrin β1 antibody (P5D2) was added at day 0 at concentrations of 0 (Ctr), 1, 2.5 and 5 µg/ml in the matrix sandwich protocol as shown in A. (D) cTnT^+^ cells measured by flow cytometry at 15 days differentiation when Cpd22 was added at day 0 at concentrations of 0 (Ctr), 1, 3 and 10 µM in the matrix sandwich protocol as shown in A. (E) Schematic for testing ILK inhibitor Cpd22 in the GiWi protocol. (F) cTnT^+^ cells measured by flow cytometry at 15 days differentiation when Cpd22 was added at day 0 at concentrations of 0 (Ctr), 1, 3 and 10 µM in the GiWi protocol as shown in E. The % of cTnT^+^ cells in each group was normalized to the control and presented as % of control. N = 3 biological replicates. Data are from DF19-9-11T iPSC line. Error bars represent SEM. *P<0.05, one-way ANOVA with post-hoc Bonferroni and Tukey test.

Integrin-linked kinase (ILK) is a serine/threonine protein kinase which interacts with the cytoplasmic domain of β1 and β3 integrins and plays crucial roles in transducing signals from ECM components and growth factors to downstream signaling components [29–31]. The kinase activity of ILK is stimulated rapidly and transiently by the engagement of integrins to FN extracellular matrix, as well as by insulin via receptor tyrosine kinases (RTKs), in a PI3K dependent manner [30]. To determine if ILK signaling is involved in the initiation of cardiac differentiation, we tested the specific ILK inhibitor, Cpd22, in the matrix sandwich protocol. Cpd22 was added at day 0 at different concentrations of 0, 1, 3 and 10 µM and cells were treated for 24 hours in the matrix sandwich protocol as shown in Fig. 6(A) Cardiac differentiation was measured by flow cytometry of the cTnT^+^ cells at 15 days of differentiation. Cpd22 showed significant inhibition of cardiac differentiation and in a concentration-dependent manner (Fig. 6D), and in multiple cell lines (Supplementary Fig. S10).

Because ILK can directly phosphorylate GSK3 and inhibit its kinase activity which modulates Wnt/beta-catenin signaling [30], we sought to confirm the role of ILK in initiation of the cardiac differentiation in a distinct hPSC-CM differentiation protocol based on biphasic modulation of Wnt signaling by small molecules. This protocol involves the activation of canonical Wnt signaling by inhibition of GSK3β to promote Brachyury^+^ precardiac mesoderm formation, followed by inhibition of Wnt to promote hPSC-CM differentiation (GiWi protocol) [32, 33]. Cpd22 was added at day 0 at different concentrations of 0, 1, 3 and 10 µM and cells were treated for 24 hours in the GiWi protocol as shown in Fig. 6E. Cardiac differentiation was measured by flow cytometry of the cTnT^+^ cells at 15 days of differentiation. Cpd22 showed significant inhibition of cardiac differentiation in a concentration-dependent manner in the GiWi protocol (Fig. 6F), similarly to the Matrix Sandwich protocol (Fig. 6C). Together these results are consistent with FN contributing to precardiac mesoderm formation and cardiac differentiation by acting via integrin β1 and ILK signaling.

## Discussion

Here we identify defined ECM proteins including LN111, LN521 and FN that can support the attachment of hPSCs and subsequent differentiation into hPSC-CMs in coordination with Activin A/BMP4/bFGF signaling. In addition, we demonstrate that initiating differentiation process with an overlay of defined ECM protein can either promote (FN) or inhibit (LN111, LN521, and COL4) cardiogenesis. Regardless of seeding substrate, cardiac differentiation starts with the generation of Brachyury^+^ cells whose survival is dependent on adequate FN in the ECM either generated endogenously or provided exogenously. FN interacts with integrins on the hPSCs including integrin β1 potentially activating ILK signaling required for the formation of Brachyury^+^ cells. These results provide new insights into defined matrices for cardiac differentiation of hPSCs and the essential role of FN in the earliest stages of differentiation.

The Activin A/BMP4/bFGF directed hPSC cardiac differentiation protocols were initially developed using the embryoid body (EB) approach starting with suspended aggregates of hPSCs [34–36]. Using the combination of Activin A/BMP4/bFGF at the initial differentiation stage in the protocol induces mesoendoderm formation consistent with known signaling pathways during embryonic development [37–40]. However, this growth factor-directed cardiac differentiation exhibited variability when applied either in the EB type protocols or monolayer-based hPSC differentiation approaches [2, 35, 41]. The lack of spatial organization and morphogen gradients in these *in vitro* differentiation approaches relative to embryonic development likely contribute to the variability. To address this variability, we developed the matrix sandwich method to promote the monolayer-based, growth factor-directed cardiac differentiation of hPSCs [2]. Although the matrix sandwich method promoted the EMT during the initial differentiation stage which is required for Brachyury^+^ mesoderm formation, the mechanism was unclear due to the complex mixture of ECM proteins in Matrigel. In the present study, we demonstrate that the accumulation and assembly of endogenously produced FN ECM induced by Matrigel overlays is a critical enabling feature of the protocol to promote formation of Brachyury^+^ precardiac mesoderm and successful cardiac differentiation.

A rapid increase in FN ECM is a well described feature of metazoan gastrulation in studies from sea urchin to mouse [42–44]. *Fn* null mouse embryos generate mesoderm but have profound heart developmental defects with early embryonic lethality [43]. FN interacts with cells by binding integrin receptors activating ILK signaling cascades including PI3K/Akt, growth factor/RTKs, GSK3 and canonical Wnt/β-catenin signaling pathways to regulate cell survival, proliferation, migration, and differentiation [30, 31]. Consistent with these developmental studies in model organisms, our data for hPSCs differentiation demonstrate that FN is critical to the genesis and survival of Brachyury^+^ precardiac mesodermal cells. FN is proposed to exert its effect by binding to integrin β1 and activating ILK signaling which can impact cell survival via PI3K/Akt signaling as well as cell transitions by inhibition of GSK3 to activate canonical Wnt signaling, which enabled the initial precardaic mesoderm formation and subsequent hPSC-CM differentiation in the Activin A/BMP4/bFGF growth factor-mediated differentiation protocol.

Over expression of ILK in epithelial cells downregulates E-cadherin, stimulates fibronectin matrix assembly and induces EMT, as well as induces the translocation of β-catenin to the nucleus, leading to the subsequent activation of the β-catenin/Tcf transcription complex, the downstream components of the Wnt signaling pathway [31, 45, 46]. We demonstrate that inhibition of ILK significantly inhibited cardiac differentiation in both the growth factor and small molecule directed protocols. The mechanism of inhibition of cardiac differentiation in the growth factor-directed, matrix sandwich protocol is likely due to the lack of Wnt/β-catenin signaling induced by ILK-GSK3β cascades signaling. On the other hand, the mechanism of inhibiting cardiac differentiation in the GiWi protocol is due to the dominant-negative effect of inhibiting ILK resulting in increased GSK3 activity [30].

The overall role of the ECM in development and promoting cell fate specification remains an important area for future investigation given the limited understanding. In addition to providing direct signaling such as FN interaction with integrins, mechanical signals from the ECM as well as the topography of the ECM directly impact the biology. For example, ECM stiffness has been suggested to be a key determinant of lineage commitment [47]. The role of the myriad of nonstructural matricellular proteins remain largely unknown. Furthermore, the ECM can provide a reservoir for more soluble signaling molecules such as TGF-β [48]. Remarkable remodeling and changes in composition of the ECM have been identified in development, but detailed understanding will require further investigation. hPSC models provide tool to investigate the role of ECM in development and differentiation, but investigations will need to consider both exogenous ECM substrates provided as well as endogenously generated ECM, e.g. the endogenously produced α-5 Laminin by undifferentiated hPSCs has been shown to promote hPSC self-renewal [49].

Overall, our study identifies several defined ECM proteins, including LN111, LN521, and FN, that can be used successfully for seeding and differentiating hPSCs to CMs. Regardless of the seeding substrate, FN in the ECM at a critical level as well as integrin β1 binding are required to stimulate the ILK signaling for successful formation of precardiac mesodermal cells required to start the cardiogenesis process whether using the Activin A/BMP4/bFGF protocol or a small molecule protocol based on biphasic Wnt modulation. Ultimately, the choice of ECM substrate used for cardiac differentiation protocols will be based not only on effective outcomes but also on the cost and availability of these defined substrates. The refinement of cardiac differentiation protocols is of growing importance as applications using hPSC-CMs expand including clinical applications.

## Materials and Methods

### Human PSC culture

Human ESC line H1 and iPSC line DF19-9-11T were used in this study. Human PSCs were cultured on Matrigel (GFR, BD Biosciences) coated 6-well plates in mTeSR1 medium. Cells were passaged using Versene solution (Gibco) every 4-5 days as previously described ([2].

### Cardiac differentiation of hPSCs

Human PSCs were dissociated with 1 ml/well Versene solution at 37°C for 5 minutes, and seeded on Matrigel, LN111 (BioLamina, 1ug/cm^2^), LN521 (BioLamina, 0.5ug/cm^2^) and FN (Corning 354008, 2.5ug/cm^2^) coated 12-well plates at the density of 6×10^5^ cells/well in mTeSR1 medium supplemented with 10 μM ROCK inhibitor (Y-27632, Tocris). The medium was changed daily. For cardiac differentiation with the matrix sandwich protocol as shown in Fig. 1A, the tested matrix proteins were overlaid on the hPSC culture after 2 (d-2) and 4 (d0) days of seeding when the monolayer of cells reached 80% and 100% confluence. The amount of matrix proteins in overlay was: 8.7ug/cm^2^ Matrigel; LN111, LN521, FN and COL4 was 0.3ug/cm^2^. For cardiac differentiation with the monolayer based protocol, the matrix overlay was not added. The cells were fed with fresh mTeSR1 medium from d-3 to d0. At d0 the medium was changed to RPMI+B27 without insulin and supplemented with growth factors. The concentration of growth factors were: Activin A (R&D, 100ng/ml), BMP4 (R&D, 2.5ng/ml) and bFGF (Waisman Biomanufacturing, UW-Madison,10ng/ml). The volume of medium was 0.8ml at d0-d1 and 1.5ml at d1-d5. The medium was changed to RPMI+B27 with insulin, and cells were fed every other day and cultured until d15 for flow cytometry analysis. For cardiac differentiation with the GiWi protocol, human PSCs were dissociated with 1 ml/well Versene solution at 37°C for 5 minutes, and seeded on Matrigel coated 12-well plates at the density of 6-10×10^5^ cells/well in mTeSR1 medium supplemented with 10 μM ROCK inhibitor (Y-27632). Cells were cultured in mTeSR1 medium with medium change daily until reached 100% confluence. At d0, the medium was changed to 1mL RPMI+B27 without insulin and supplemented with 12 µM CHIR99021 (Tocris), and changed to 2mL RPMI+B27 without insulin after 24h. 72 h after addition of CHIR99021, a combined medium was prepared by collecting 1mL of medium from the wells and mixing with same volume of fresh RPMI+B27 without insulin medium and supplemented with 5 µM IWP2 (Tocris). The medium was changed to 2mL RPMI+B27 without insulin at d5, and to RPMI+B27 with insulin starting from d7. Cells were fed every other day and cultured until d15 for flow cytometry analysis.

### Generation of inducible *FN1* knockdown hPSC clones

Human PSCs maintained in mTeSR1 medium were used for lentivirus transfection. The cloning strategy was schemed in Supplementary Fig. S12.

#### Production of lentivirus particles

HEK 293 TN cells (SBI) were used for lentivirus production. Briefly, 4.5 x 10^6^ cells were plated in a 10cm dish on the day before transfections. Lipofectamine 2000 (Invitrogen) was used for transfections (1:2 ratio). Transfection conditions used were – 7ug lentivirus plasmid, 10ug psPAX2 (Addgene plasmid #12260 - packaging), 5ug pMD2.G (Addgene plasmid #12259 - envelope). Transfection media was incubated for 15-16 hours, after which it was replaced with 5mls of mTeSR1medium. Lentivirus supernatant was collected after 48-52 hours, filtered (0.45 uM – Millipore) and frozen down at -80°C. Lentivirus supernatants were thawed in 37°C water bath immediately before infection.

#### hPSCs transfection and neomycin selection

Human PSCs at the 80% confluency after split were incubated with the lentivirus in mTeSR1 medium in the presence of 8ug/ml Polybrene (Sigma) for 24-42 hours. After lentivirus transfection, hPSCs were washed with PBS and recovered in mTeSR1 medium for 2-3 days with medium change daily, cells were split once during this recovery. The concentrations of G418 (Life technologies) for neo-resistant selection for hPSC lines were determined by the kill curve (Supplementary Fig. S13). G418 (100mg/ml) were diluted in mTeSR1 medium with the final concentrations of 100ug/ml for H1 ES cells and 75ug/ml for DF19-9-11T, and hPSCs were under neomycin selectin up to 11 days before single cell isolation and cloning.

#### Single cell isolation and cloning

Human PSCs resistant to neomycin were incubated with Accutase (Gibco) in 37°C for 5-10 minutes, and resuspended in mTeSR1 medium to examine under microscope for single cells. If cells were still clustered, cells were spun down and washed with PBS, and underwent Accutase treatment for another time as above. Single cells were plated in 6-well Matrigel coated plate at the very low density of 50-100 cells/cm^2^ with 10uM Rock inhibitor (Y-27632). Single cells were growing in mTeSR1 medium for 4-7 days, clones were picked up by P200 mircropipette tips and transferred to microtubes with 30ul Versene solution (pre-warmed at room temperature), and incubated at 37°C for 5 minutes. Cells were pipette up and down several times and equally dispersed in two wells of Matrigel coated 24-well plates in mTeSR1 medium.

#### Doxycycline induction

To induce *FN1* shRNA and mCherry expression, doxycycline (Sigma, 8ug/ml) in mTeSR1 medium was added to the hPSC clones culture in the 12-well plate for 2-3 days. mCherry expression were examined with EVOS microscope (Life Technologies); *FN1* expression were examined by quantitative RT-PCR.

#### Clones expansion and cryopreservation

Selected *FN1* knockdown positive clones were expanded in mTeSR1 medium in 6-well culture. Cells were cryopreserved in 90%FBS, 10%DMSO and 10uM ROCK inhibitor in liquid nitrogen freezer (LABS20K, Taylor-Wharton).

### Flow cytometry

Cells were detached from cell culture plates by incubation with 0.25% trypsin-EDTA (Invitrogen) plus 2% chick serum (Sigma) for 5 minutes at 37°C. Cells were vortexed to disrupt the aggregates followed by neutralization by adding equal volume of EB20 medium.[50] Cells were fixed in 1% paraformaldehyde at 37°C water bath for 10 minutes in the dark, permeabilized in ice-cold 90% methanol for 30 minutes on ice. Cells were washed once in FACS buffer (PBS without Ca/Mg^2+^, 0.5% BSA, 0.1% NaN3) plus 0.1% Triton, centrifuged, and supernatant discarded leaving about 50 μl. For labeling cTnT, the primary antibody was diluted in 50 μl/sample FACS buffer plus 0.1% Triton and aliquoted to each sample for a total sample volume of 100 μl. Samples were incubated with the primary antibodies overnight at 4°C. For co-labeling of Brachyury and Sox17, the cells were incubated with the conjugated antibodies for half an hour at room temperature in dark. Cells were washed once with 3 ml FACS buffer plus 0.1% Triton and resuspended in 300 – 500 μl FACS buffer plus Triton for analysis. Please refer to Supplementary Material for detail of the primary antibodies. For the secondary antibody labeling for cTnT, cells were washed once with 3 ml FACS buffer plus 0.1% Triton after the primary antibody labeling, centrifuged, and supernatant discarded leaving ∼ 50 μl. Secondary antibody specific to the primary IgG isotype was diluted in FACS buffer plus Triton in a final sample volume of 100 μl at 1:1000 dilution. Samples were incubated for 30 minutes in the dark at room temperature, washed in FACS buffer plus Triton and resuspended in 300 – 500 μl FACS buffer plus Triton for analysis. Data were collected on a FACSCaliber flow cytometer (Beckton Dickinson) and analyzed using FlowJo.

### Immunocytochemistry

For imaging with EVOS microscope (Life technologies), hPSCs were cultured either in 6-well plates, or seeded and differentiated in 12-well plates, coated with Matrigel or the ECM proteins. For imaging with confocal microscope (Leica), cells were seeded or plated on glass coverslips coated with Matrigel or the ECM proteins. Cells were fixed with 4% paraformaldehyde for 15 minutes at room temperature for labeling with different markers. For intracellular markers, cells were permeabilized in 0.2% Triton X-100 (Sigma) for 1 hour at room temperature. For labeling the ECM proteins, cells were not permeabilized. After fix and permeablization, samples were blocked with 5% non-fat dry milk (Bio-Rad) in 0.2% Triton X-100 solution and incubated for 2 hours at room temperature on a rotator followed by two washes with PBS. Primary antibodies (please refer to Supplementary Material for detail of the primary antibodies) were added in 1% BSA in PBS solution with or without 0.1% Triton X-100 depends on the markers to label, and incubated overnight at 4°C. Samples were washed with 0.2% Tween 20 in PBS twice and 1X PBS twice. Secondary antibody specific to the primary IgG isotype were diluted (1:1000) in the same solution as the primary antibodies and incubated at room temperature for 1.5 hours in dark on a rotator. Samples were washed with 0.2% Tween 20 in PBS twice and 1X PBS twice. Nuclei were labeled with Hoechst or DAPI. Confocal images were analyzed with the Leica LAS AF Lite software.

### Quantification of fibronectin immuno-fluorescence by ImageJ analysis

The fibronectin immuno-fluorescence images of d-3, d-2, d-1, d0 matrix sandwich culture, and d0 monolayer culture were evenly split into 4 squares with online image splitting tool (https://www.imgonline.com.ua/eng/cut-photo-into-pieces.php), and imported to ImageJ Fiji software. The control images without primary antibody labeling were also imported and analyzed in the same way by ImageJ Fiji software. The gray value was used to represent the intensity of fluorescence, ranging from 0 (black, low intensity) to 256 (white, high intensity). For mean and standard deviation of gray values, the entire area of each image was selected, and the values were retrieved by selecting Menu bar > Analyze > Measure. The mean gray value was generated from ImageJ by summing each pixel’s gray value divided by the number of pixels in the selected area. The final mean fluorescence value was obtained by subtracting the mean gray value of the control image from fibronectin immuno-labeled images. The mean of fluorescence intensity of each quartiles from each biological replicate in each group were represented in the box plots using Origin v9.

### RT-PCR and quantitative RT-PCR

Cell samples were collected using 0.25% trypsin-EDTA (Invitrogen) to remove the cells from cell culture plates. Total RNA was purified using QIAGEN RNeasy® Mini kit. Possible genomic DNA contamination was removed by RNase-Free DNase Set (QIAGEN) with the RNeasy columns, or by DNase I (Invitrogen) treatment for 15 minutes at room temperature. 500 ng of total RNA was used for Oligo(dT)20 – primed reverse transcription using SuperScript™ III First-Strand Synthesis System (Invitrogen). Quantitative RT-PCR was performed using Taqman PCR Master Mix and Gene Expression Assays (Applied Biosystems, Supplementary Table 1) in triplicate for each sample and each gene. 0.5 μl of cDNA from RT reaction was added as template for each Q-PCR reaction. The expression of genes of interest was normalized to that of *GAPDH*. RT-PCR was carried out using Platinum™ Taq DNA Polymerase (Invitrogen) or Gotaq Master Mix (Promega) and then subjected to 2% agarose gel electrophoresis. PCR conditions included denaturation at 94°C for 30 seconds, annealing at 60°C for 30 seconds, and extension at 72°C for 1 minute, for 35 cycles, with 72°C extension for 7 minutes at the end. *ACTB* (β-actin) was used as an endogenous control.

### Antibody blocking and small molecule inhibition

Blocking antibodies without sodium azide were used for the blocking experiments. Blocking antibody for integrin β1, P5D2 (Developmental Studies Hybridoma Bank), and for integrin α5, P1D6 (Developmental Studies Hybridoma Bank), and ILK inhibitor Cpd22 (EMD Millipore) were diluted in the media to make the final concentrations and added to the culture at the time points as shown in Fig. 6.

### Statistics

Data are presented as mean ± standard error of the mean (SEM). Technical and biological replicates are as indicated for each dataset. For datasets with normal distributions, statistical significance was determined by Student’s t-test for two groups or one-way ANOVA for multiple groups with post-hoc test using Bonferroni and Tukey methods. Statistical analysis was performed using Origin, v9, P < 0.05 was considered statistically significant.

## ACKNOWLEDGMENTS

The authors thank Dr. Deane Mosher for the consultation of fibronectin biology and Dr. Douglas Annis to assist with antibody epitope mapping. The authors also thank Ms. Yukun Li to assist with the image analysis using ImageJ. The work was funded by NIH R01 HL129798 (T.J.K.) and NIH U01HL134764 (T.J.K.).

## AUTHOR CONTRIBUTIONS

TJK and JZ conceived the study and wrote the manuscript. JZ designed and carried out the experiments, collected and analyzed the data. RT did most of the cell culture and the qRT-PCR. JLC did the RT-PCR. PAL did the FN knockdown construct. YM did the western blot. SPP contributed funding support and writing/editing the manuscript.

---

**Supplementary Fig. S1.**
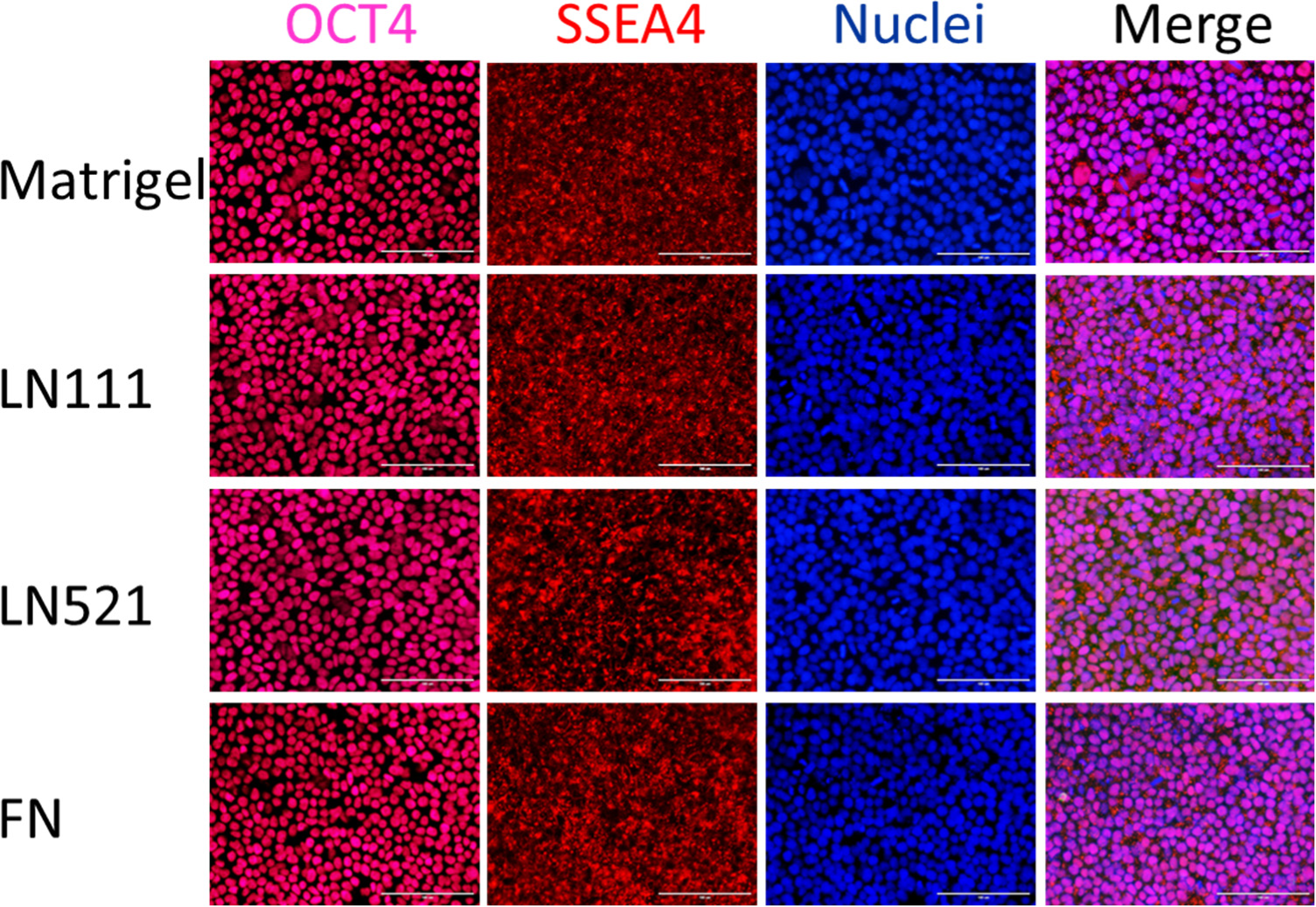
Fluorescence images of H1 ESCs growing on the ECM of Matrigel, LN111, LN521 and FN as confluent monolayer and immunolabeled with antibodies against OCT4 and SSEA4. Scale bar is 100 µm.

**Supplementary Fig. S2.**
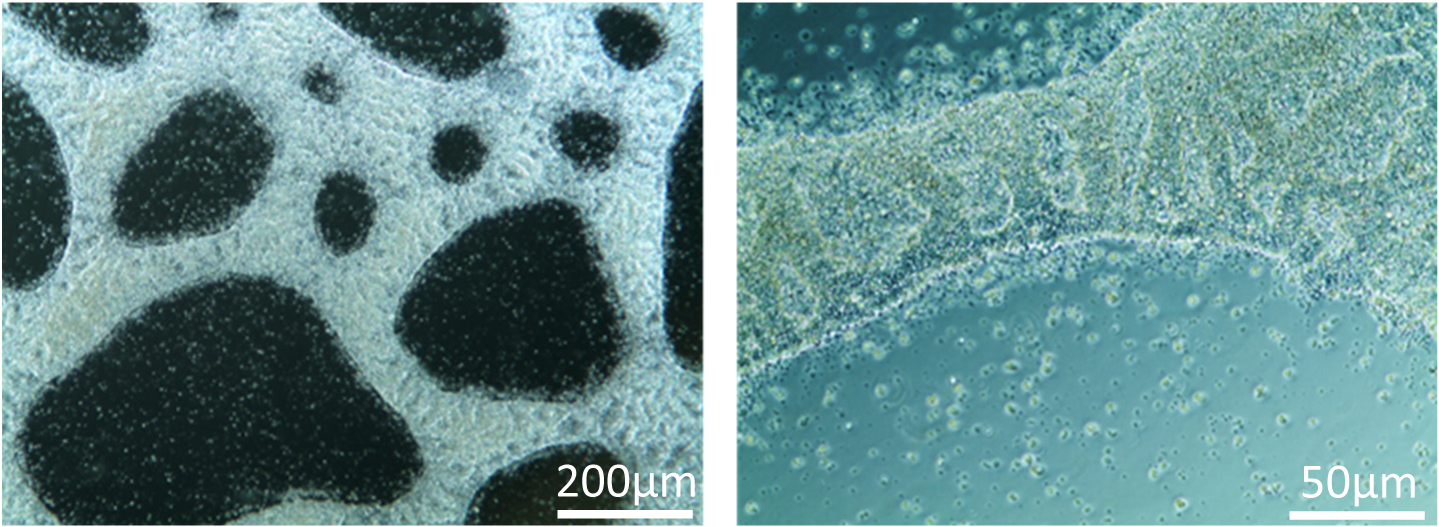
Morphology of DF19-9-11 iPSCs growing on human collagen IV coated surface. The iPSCs growing for 5 days on collagen IV coated surface do not form confluent monolayer.

**Supplementary Fig. S3.**
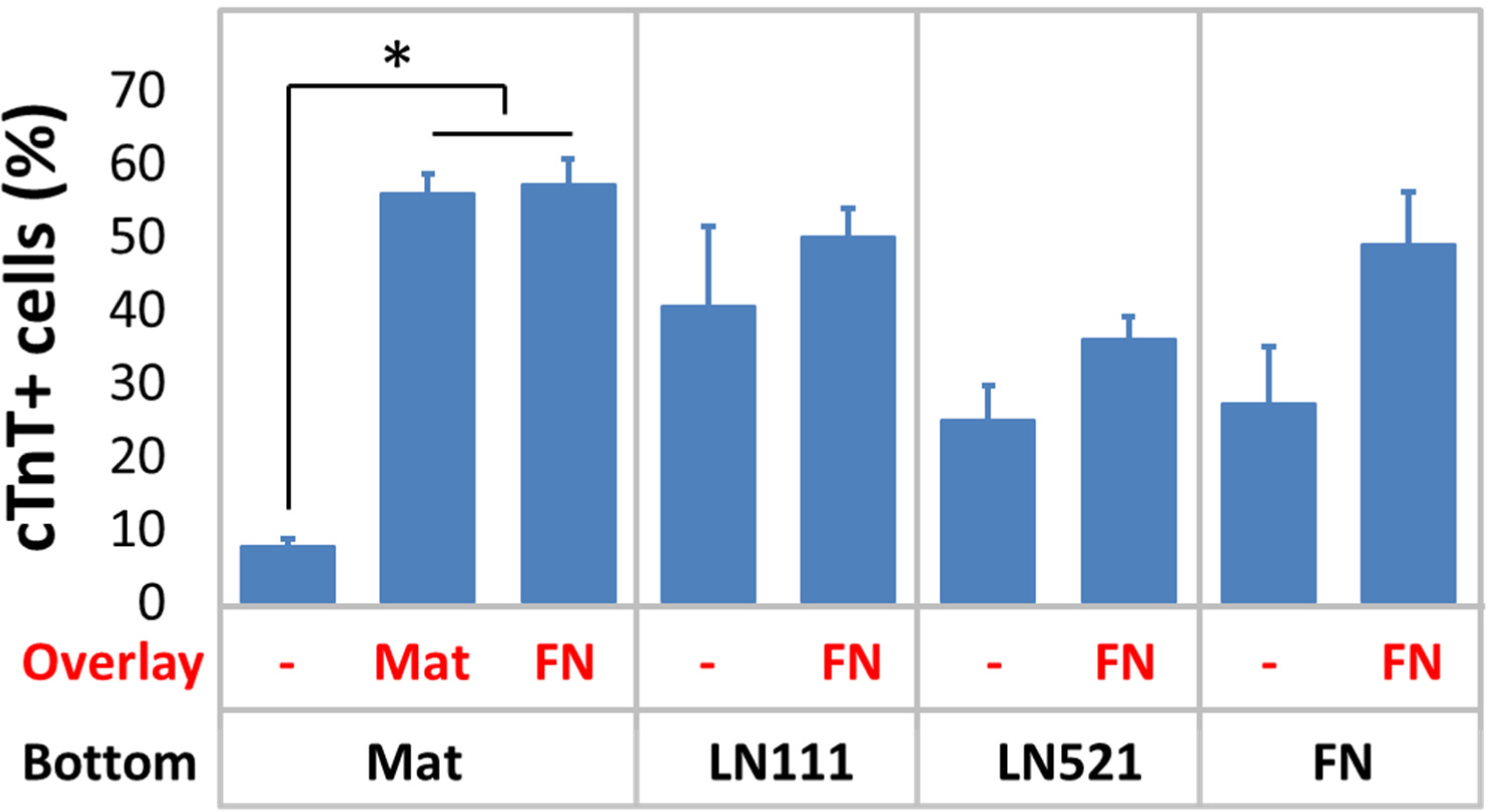
cTnT^+^ cells measured by flow cytometry at 15 days of differentiation of H1 ESCs on different ECM proteins as substrate (bottom) and overlay (overlay) using the matrix sandwich protocol. N ≥ 3 biological replicates. *P<0.05, one-way ANOVA with post-hoc Bonferroni and Tukey test.

**Supplementary Fig. S4.**
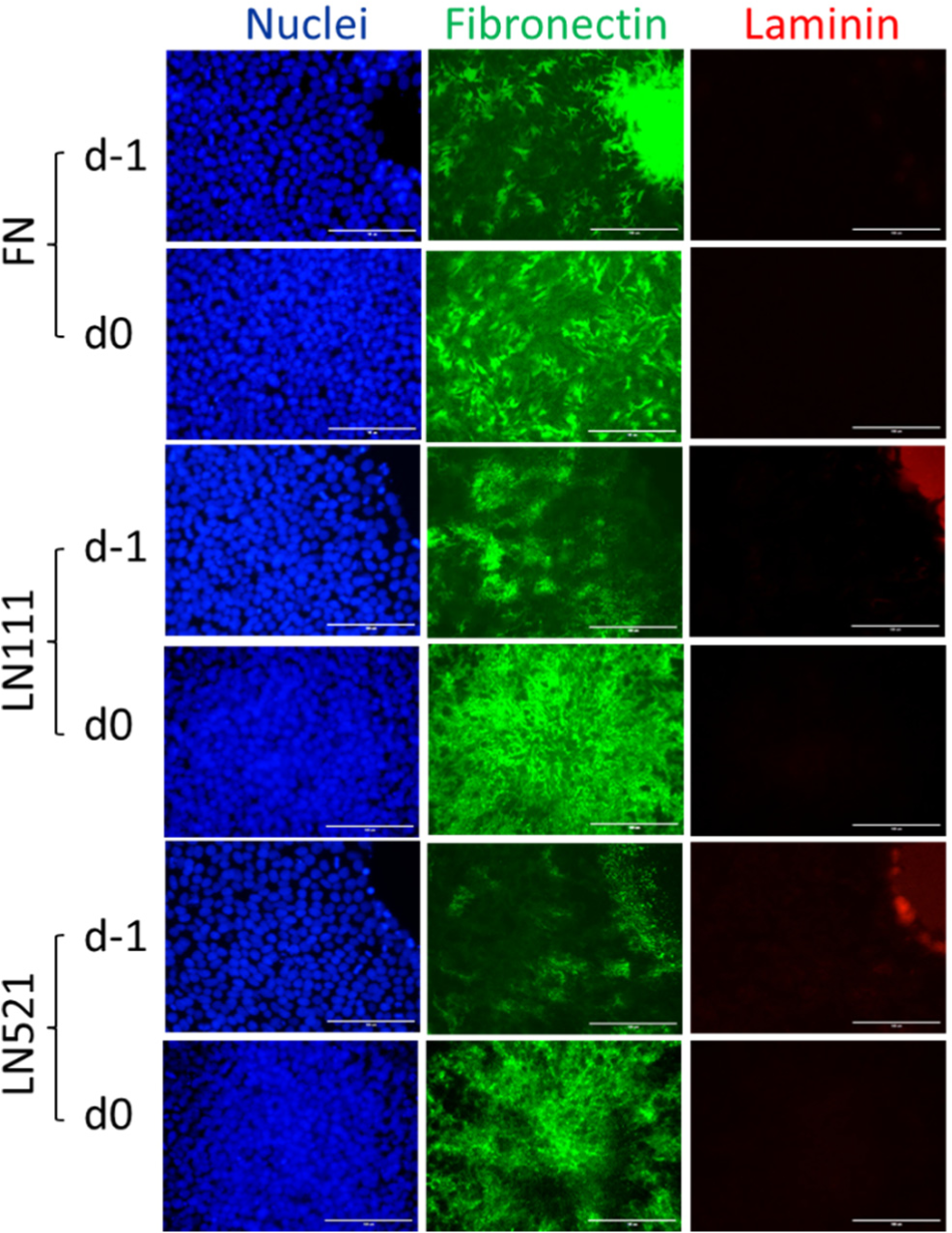
Fluorescence images of hPSCs growing on FN, LN111 and LN521 coated surface immunolabeled with antibodies against fibronectin and laminin. Scale bar is 100 µm.

**Supplementary Fig. S5.**
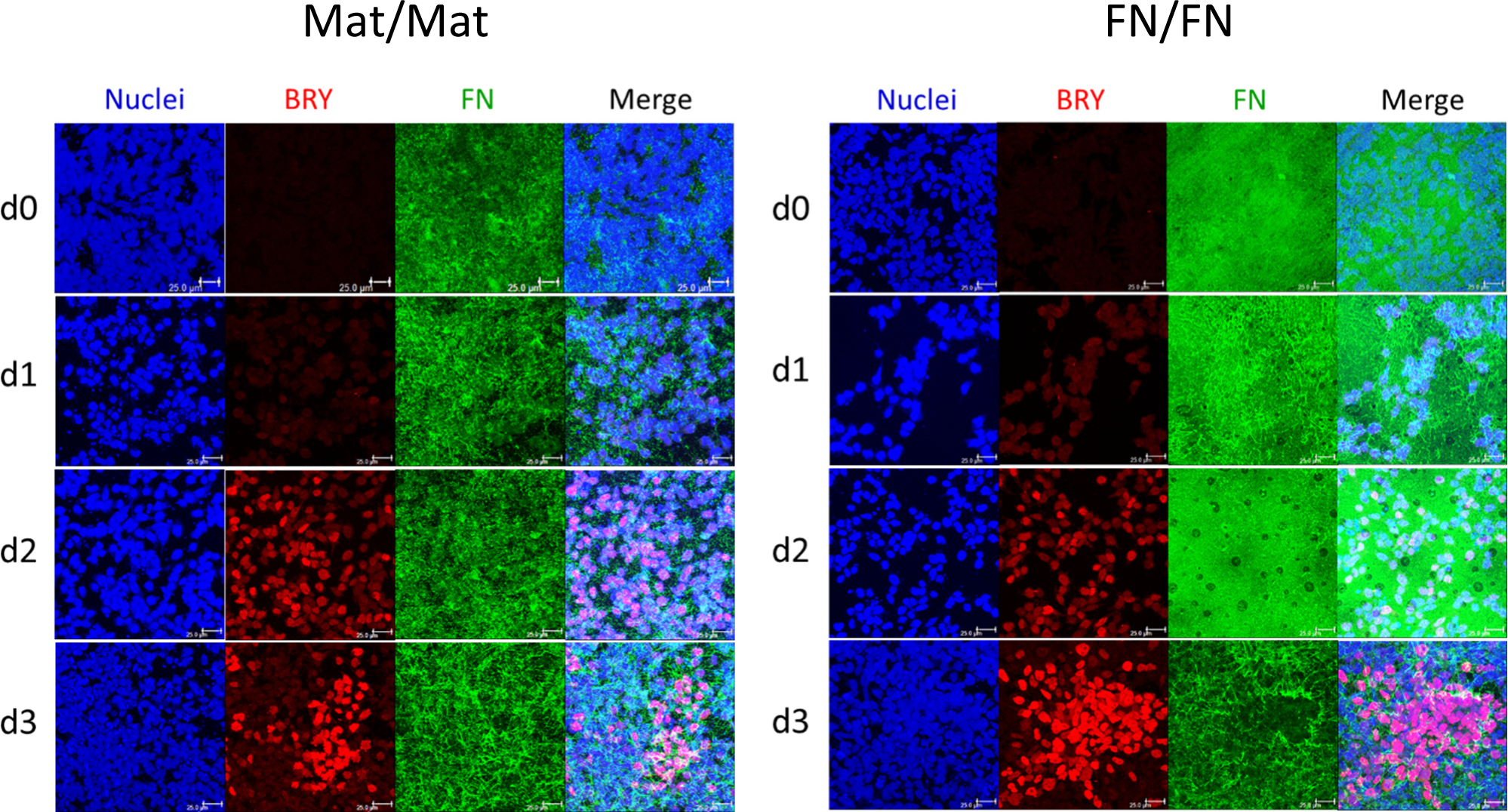
Maximum projection view of the confocal z-scan of DF19-9-11T iPSCs at day 0-3 of the cardiac differentiation using the Matrigel/Matrigel (Mat/Mat) and FN/FN matrix sandwich protocol. The cells were co-labeled with antibodies against brachyury (BRY) and fibronectin (FN). Scale bar is 25 µm.

**Supplementary Fig. S6.**
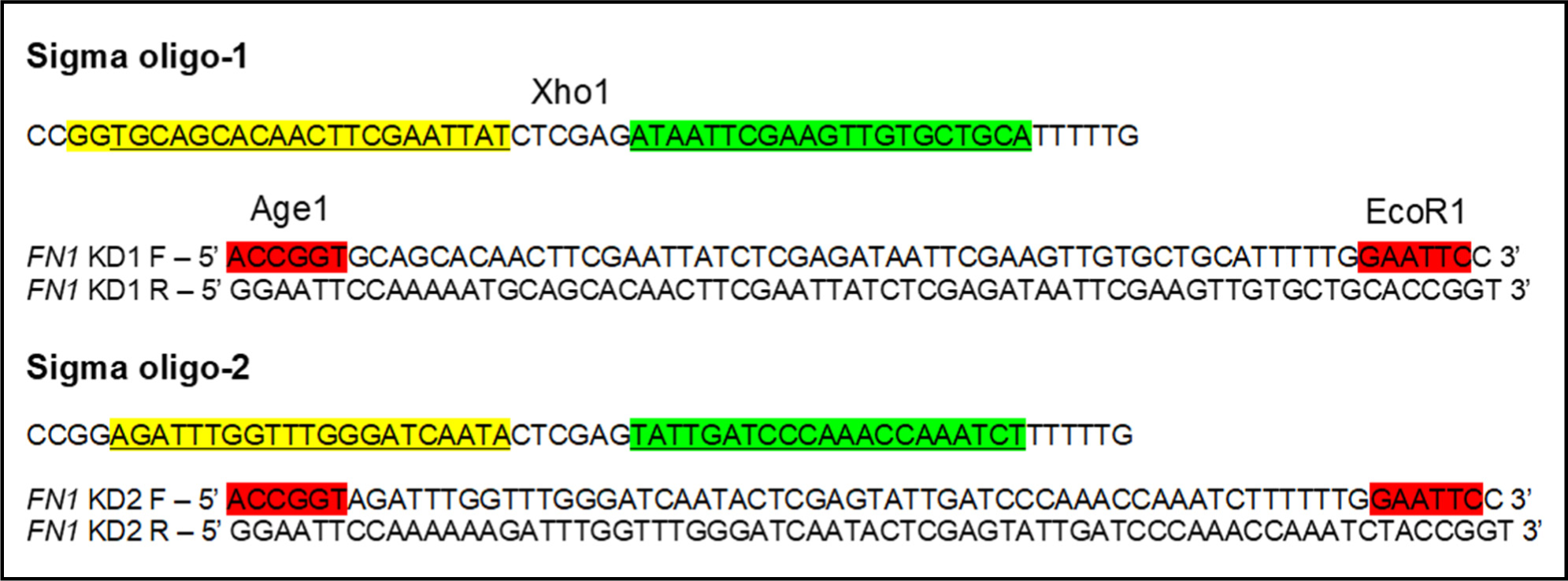
The *FN1* shRNA oligos. Yellow, CDS where shRNA binds; <u>Green</u>, palindrome. Oligo-1 binds in exon 8 (95% KD); Oligo-2 binds in 3’ UTR (93% KD).

**Supplementary Fig. S7.**
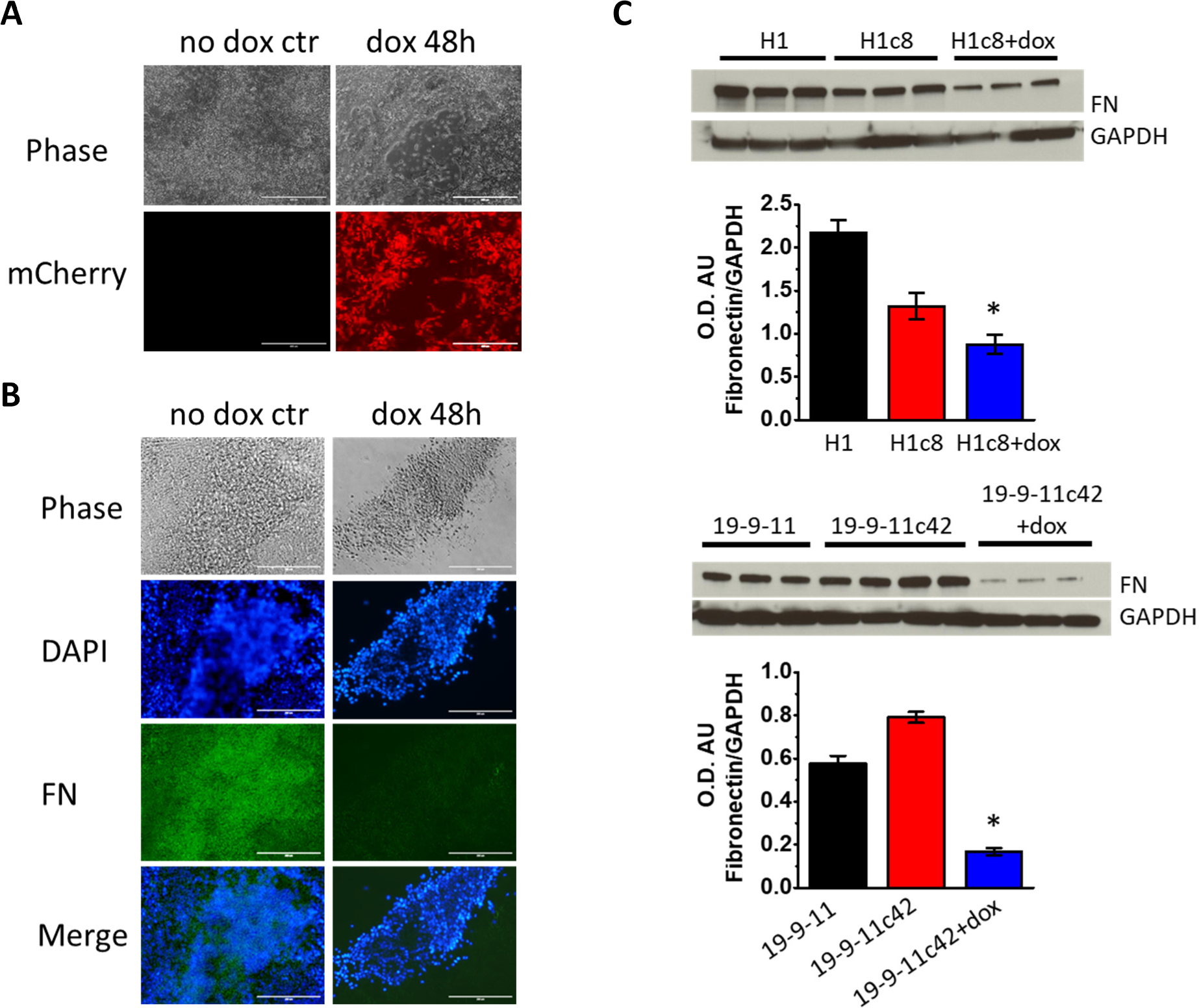
Characterization of hPSC *FN1* knockdown clones. **A** Doxycycline induced mCherry expression in the hPSC culture of H1 *FN1* knockdown clone. Scale bar is 200 µm. **B** Phase contrast and immuno-fluorescence images of FN expression in doxycycline induced hPSC culture of H1 *FN1* knockdown clone. Scale bar is 200 µm. **C** Quantatitave western blot of FN expression in hPSC culture of H1 and 19-9-11 *FN1* knockdown clones. N = 3 biological replicates. *P<0.05, one-way ANOVA with post-hoc Bonferroni and Tukey test.

**Supplementary Fig. S8.**
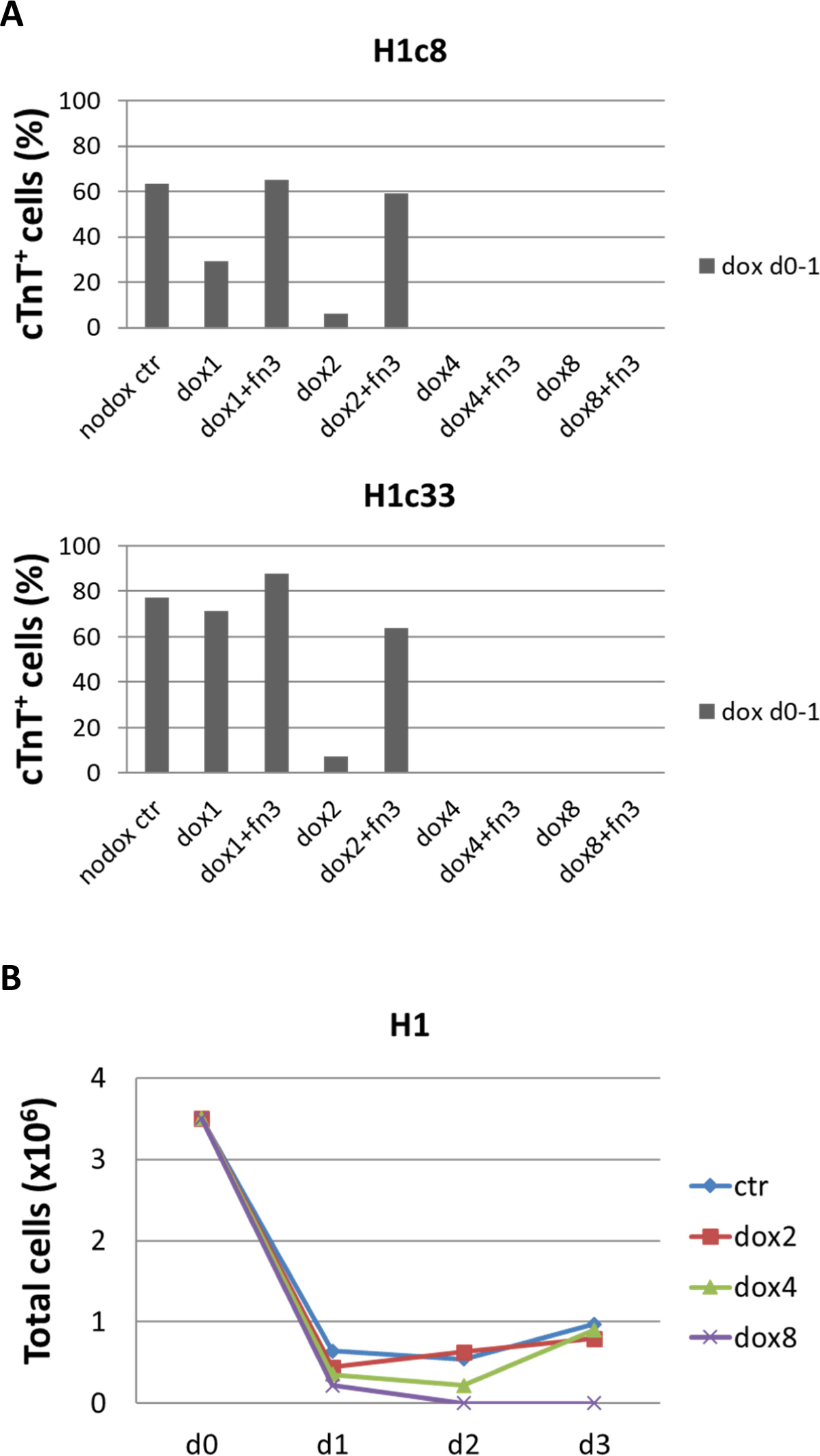
FN knockdown at day 0-1 of the cardiac differentiation is doxycycline concentration dependent. **A** cTnT^+^ cells measured by flow cytometry at 15 days of differentiation of the *FN1* knockdown clones (H1c8 and H1c34) showing the effect of FN knockdown at day 0-1 at different concentrations of dox and the effect of adding exogenous FN at each concentration of dox. Dox concentration higher than 4ug/ml caused significant cell death, and cardiac differentiation could not be rescued by exogenous FN. **B** The total cells of the regular H1 line in the same cardiac differentiation at day 0-3 when dox was added at day 0-1. Dox concentrations of 0 (nodox ctr), 1 (dox1), 2 (dox2), 4 (dox4) and 8 (dox8) ug/ml were tested. +fn indicates exogenous FN (3ug/cm^2^) was added at day 0- 1 for each concentration of dox group.

**Supplementary Fig. S9.**
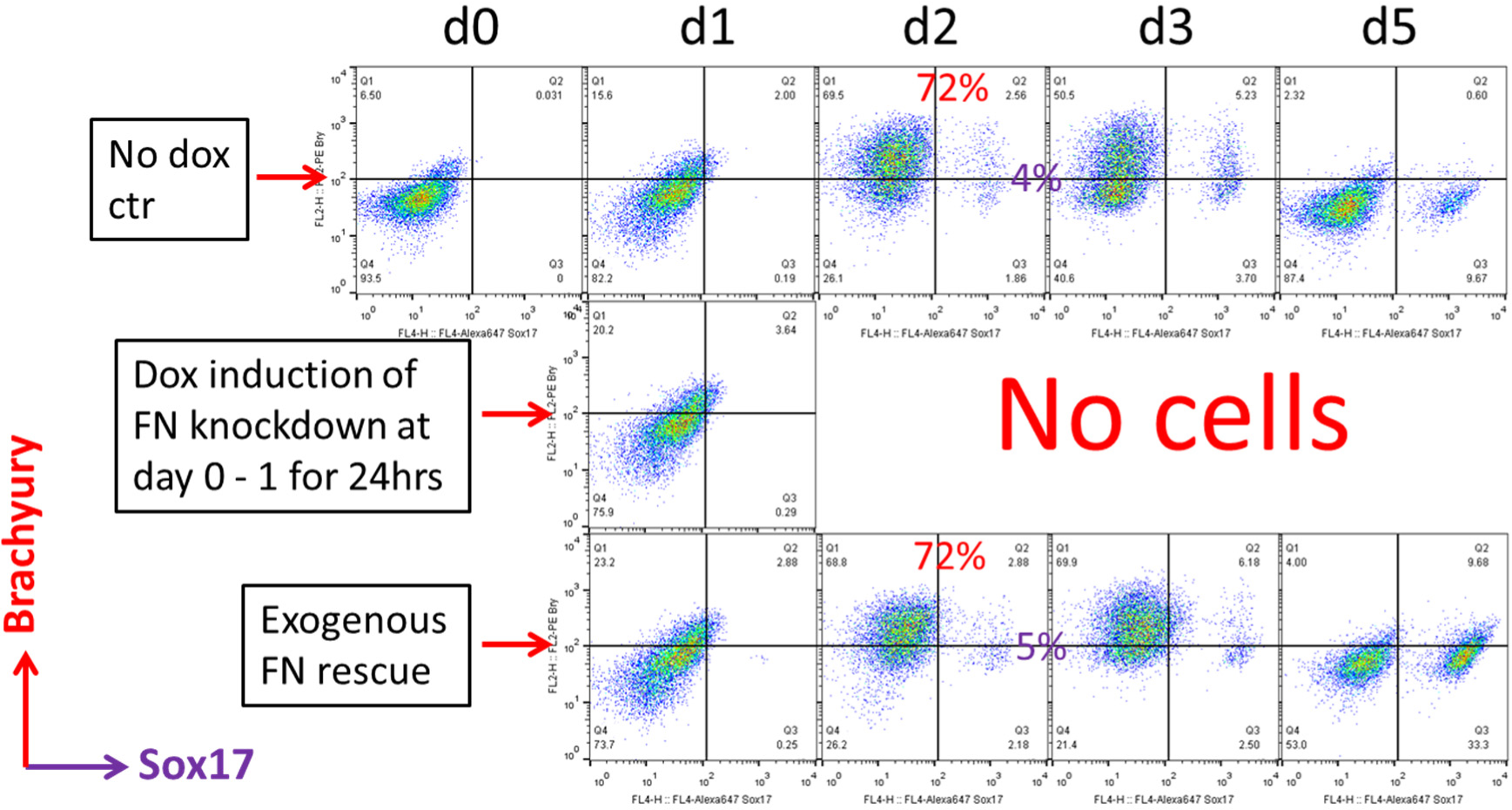
Flow cytometry of co-labeling the cells with Brachyury and Sox17 antibodies of the 19-9-11 FN knockdown clone in the cardiac differentiation time course of day 0-5 at the no dox control, dox induction at day 0-1 and dox induction at day 0-1 with adding exogenous FN conditions.

**Supplementary Fig. S10.**
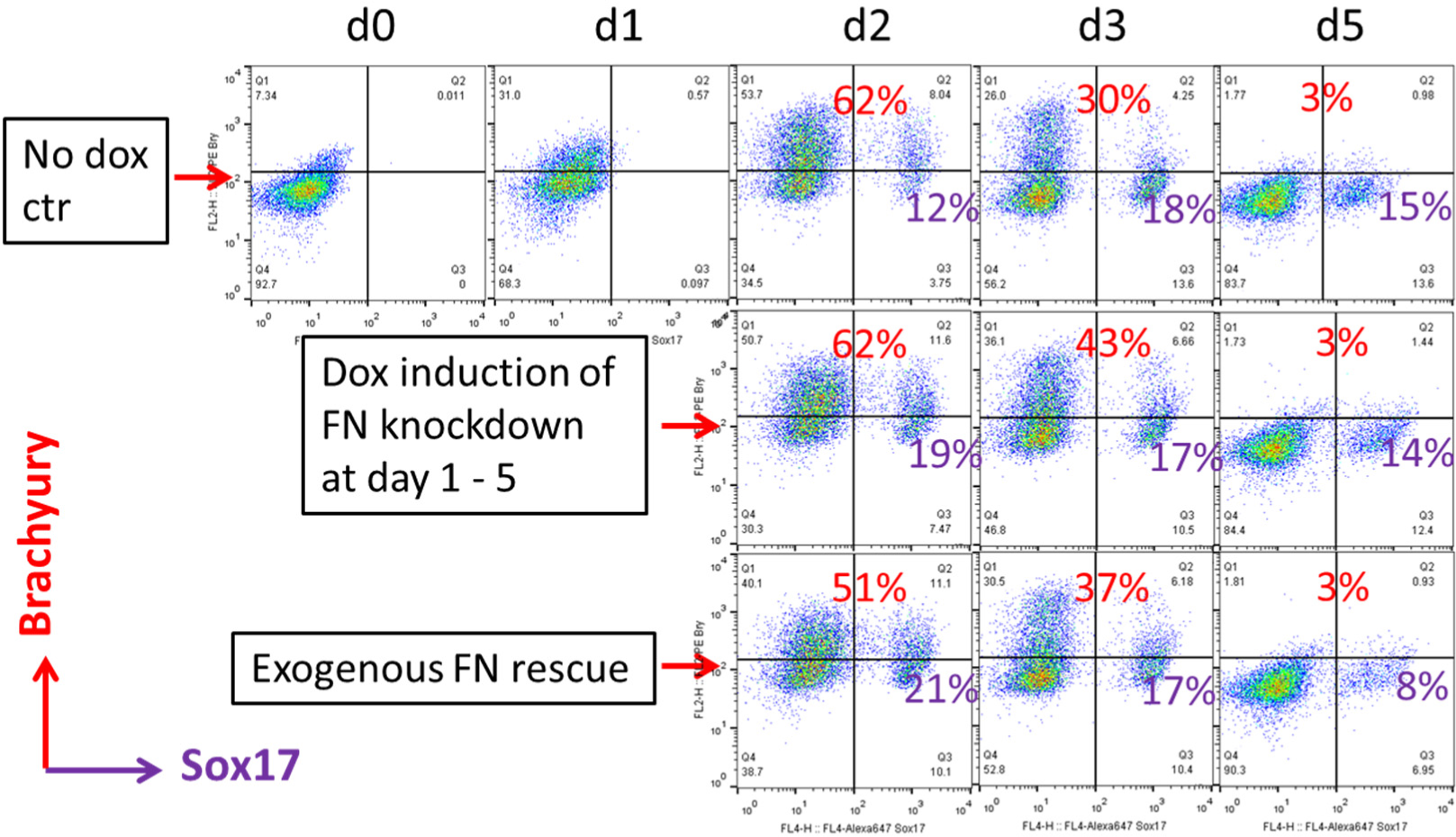
Flow cytometry of co-labeling the cells with Brachyury and Sox17 antibodies of the H1 FN knockdown clone in the cardiac differentiation time course of day 0- 5 at the no dox control, dox induction at day 1-5 and dox induction at day 1-5 with adding exogenous FN conditions.

**Supplementary Fig. S11.**
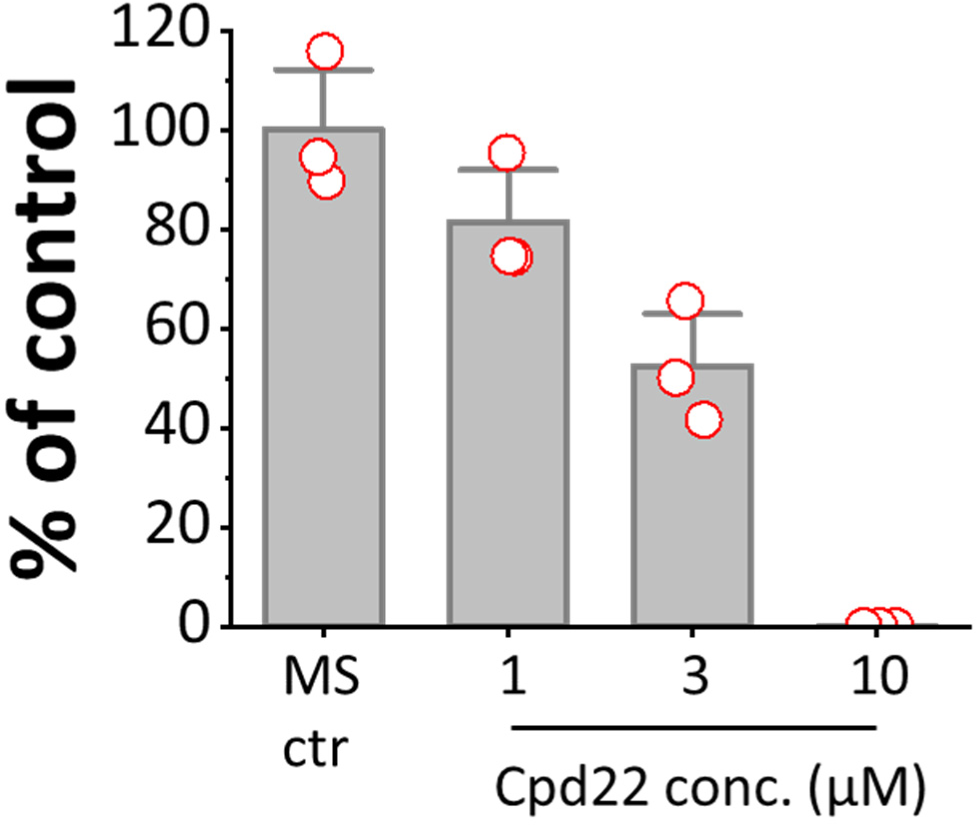
The ILK inhibitor, Cpd22 was added at day 0 at concentrations of 0, 1, 3 and 10 uM in the matrix sandwich protocol. Cardiac differentiation was measured by flow cytometry of the cTnT^+^ cells at 15 days differentiation. The cTnT^+^ cells in each group was normalized to the control and shown as % of control on the plot. N = 3 biological replicates. Data are form H1 ESC line.

**Supplementary Fig. S12.**
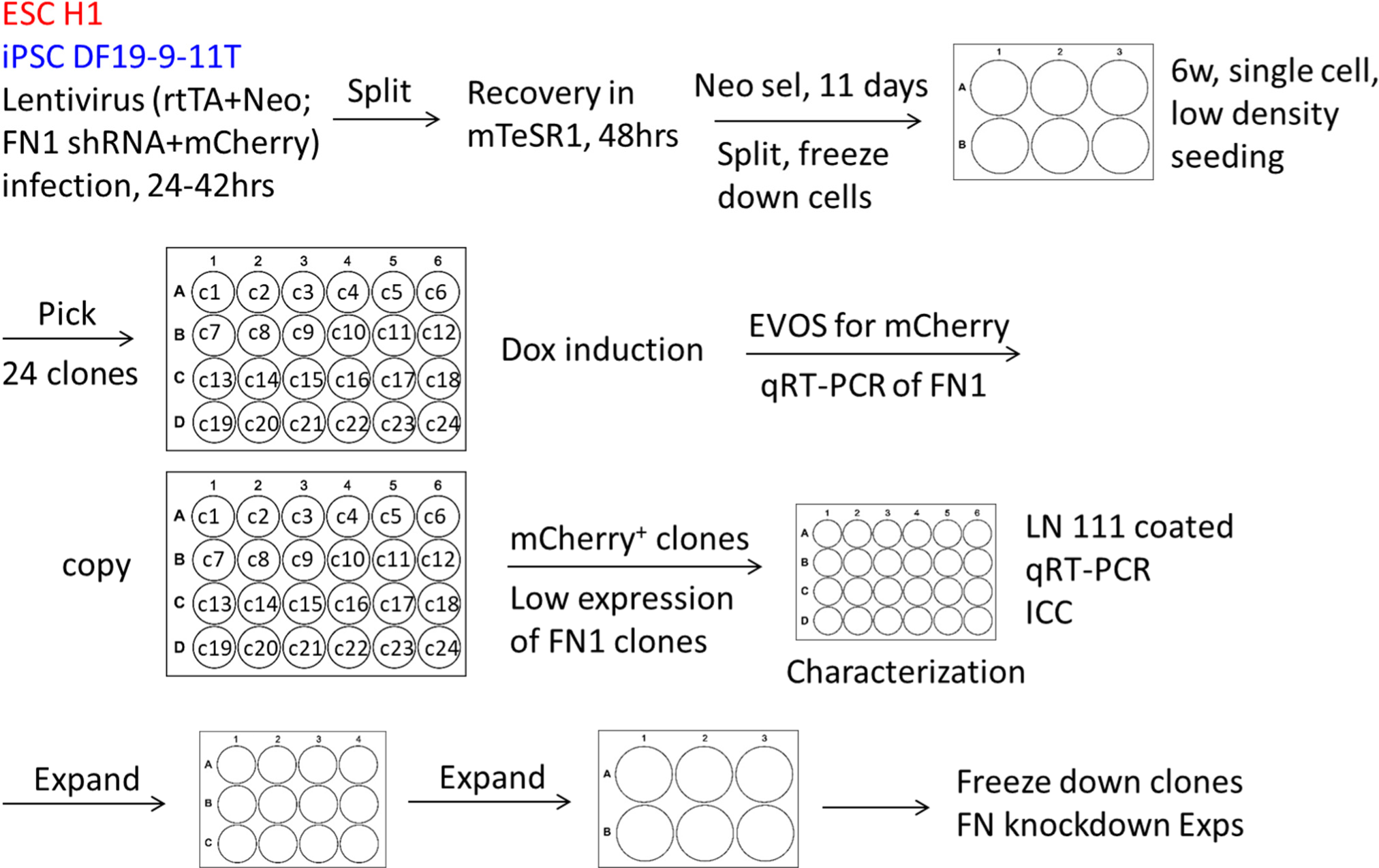
The cloning strategy in generation of inducible *FN1* knockdown hPSC clones

**Supplementary Fig. S13.**
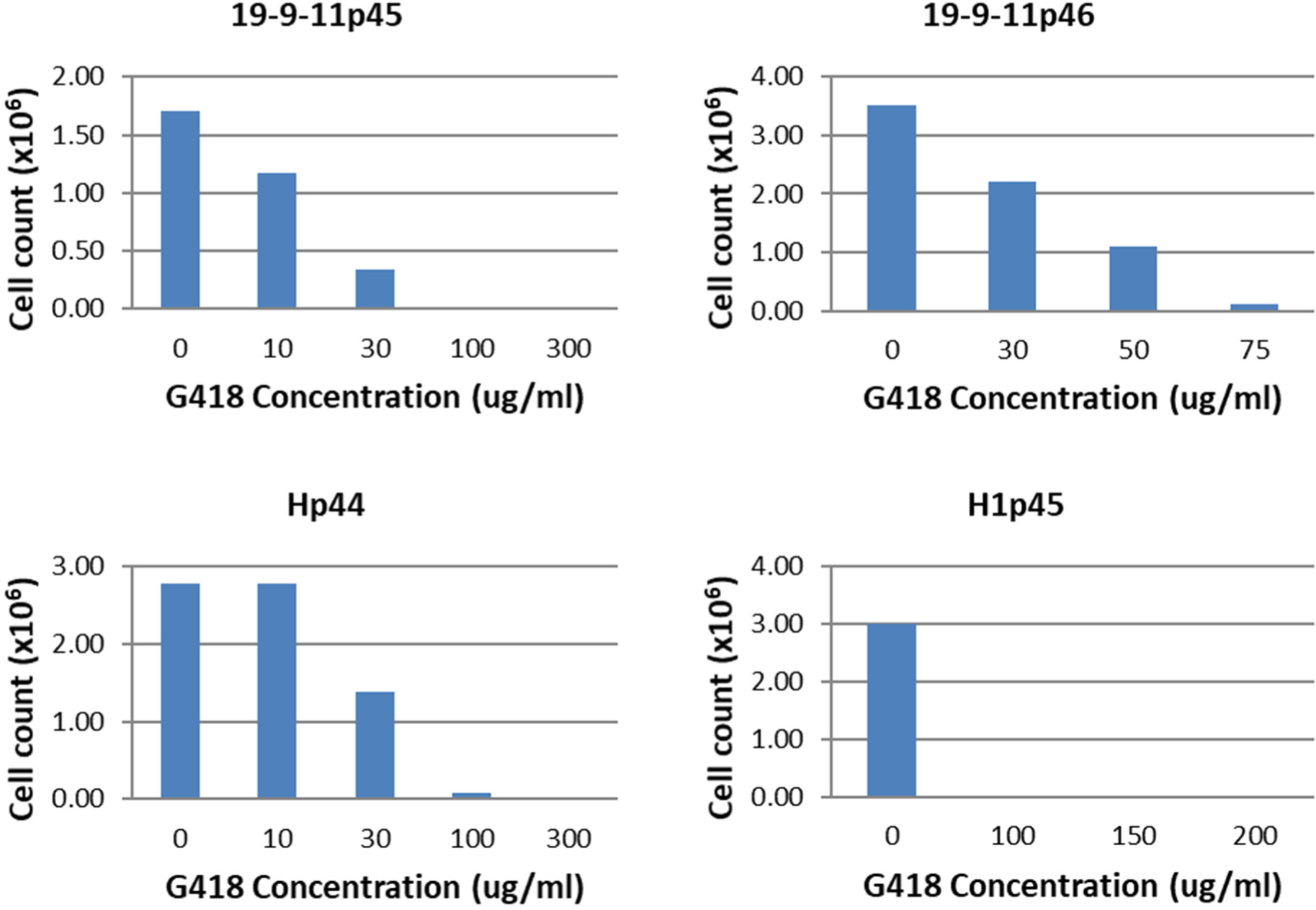
The concentrations of G418 test for neo-resistant selection for hPSC lines.

**Supplementary Table 1.** Primary antibodies used in flow cytometry (FC) and immunocytochemistry (ICC)

|  |  |  |  |  |
| --- | --- | --- | --- | --- |
| Human/Mouse Brachyury PE-conjugated | R&D (IC2085P) | Goat | IgG | 10ul/1x10<sup>6</sup> cells (FC) |
| Alexa Fluor 647 Mouse Anti-Human Sox17 (Clone P7-969) | BD (562594) | Mouse | IgG<sub>1</sub> | 5ul/1x10<sup>6</sup> cells (FC) |
| Human/Mouse Brachyury Antibody | R&D (AF2085) | Goat | IgG | 5ug/ml (ICC) |
| Troponin T, Cardiac Isoform Ab-1 (Clone 13-11) | ThermoFisher Scientific (#MS-295-P) | Mouse | IgG<sub>1</sub> | 1:200 dilution (FC, ICC) |
| Purified Mouse Anti-Fibronectin (Clone 10) | BD (61007) | Mouse | IgG<sub>1</sub> | 1:100 dilution (ICC) |
| Anti-Laminin | Sigma-Aldrich (L9393) | Rabbit | IgG | 1:500 dilution (ICC) |
| Oct-3/4 (C-10) | Santa Cruz (sc-5279) | Mouse | IgG<sub>2b</sub> | 1:100 dilution (ICC) |
| Anti-SSEA4 antibody [MC813-70] | Abcam (ab16287) | Mouse | IgG<sub>3</sub> | 1:200 dilution (ICC) |

**Supplementary Table 2.** Primers for quantitative RT-PCR

| **Genes** | **TaqMan® Gene Expression Assay ID** |
| --- | --- |
| GAPDH | Hs99999905_m1 |
| SNAIL1 | Hs00195591_m1 |
| SNAIL2 | Hs00950344_m1 |
| TWIST1 | Hs00361186_m1 |
| VIM | Hs00185584_m1 |
| FN1 | Hs01549976_m1 |
| CDH1 | Hs01023894_m1 |
| CDH2 | Hs00983056_m1 |
| GSC | Hs00418279_m1 |
| MIXL1 | Hs00430824_g1 |
| SOX17 | Hs00751752_s1 |
| TBXT | Hs00610080_m1 |
| MESP1 | Hs00251489_m1 |
| ISL1 | Hs00158126_m1 |
| NKX2-5 | Hs00231763_m1 |
| GATA4 | Hs00171403_m1 |

## REFERENCES

[1] D.A.C. Walma, K.M. Yamada, The extracellular matrix in development, Development, 147 (2020).

[2] J. Zhang, M. Klos, G.F. Wilson, A.M. Herman, X. Lian, K.K. Raval, M.R. Barron, L. Hou, A.G. Soerens, J. Yu, S.P. Palecek, G.E. Lyons, J.A. Thomson, T.J. Herron, J. Jalife, T.J. Kamp, Extracellular matrix promotes highly efficient cardiac differentiation of human pluripotent stem cells: the matrix sandwich method, Circ Res, 111 (2012) 1125–1136.

[3] M.A. Nieto, R.Y. Huang, R.A. Jackson, J.P. Thiery, Emt: 2016, Cell, 166 (2016) 21–45.

[4] J.W. Lambshead, L. Meagher, C. O’Brien, A.L. Laslett, Defining synthetic surfaces for human pluripotent stem cell culture, Cell Regen, 2 (2013) 7.

[5] S. Rodin, L. Antonsson, C. Niaudet, O.E. Simonson, E. Salmela, E.M. Hansson, A. Domogatskaya, Z. Xiao, P. Damdimopoulou, M. Sheikhi, J. Inzunza, A.S. Nilsson, D. Baker, R. Kuiper, Y. Sun, E. Blennow, M. Nordenskjold, K.H. Grinnemo, J. Kere, C. Betsholtz, O. Hovatta, K. Tryggvason, Clonal culturing of human embryonic stem cells on laminin-521/E-cadherin matrix in defined and xeno-free environment, Nature communications, 5 (2014) 3195.

[6] L. Yap, J.W. Wang, A. Moreno-Moral, L.Y. Chong, Y. Sun, N. Harmston, X. Wang, S.Y. Chong, K. Vanezis, M.K. Ohman, H. Wei, R. Bunte, S. Gosh, S. Cook, O. Hovatta, D.P.V. de Kleijn, E. Petretto, K. Tryggvason, In Vivo Generation of Post-infarct Human Cardiac Muscle by Laminin-Promoted Cardiovascular Progenitors, Cell reports, 26 (2019) 3231–3245 e3239.

[7] J.P. Jung, D. Hu, I.J. Domian, B.M. Ogle, An integrated statistical model for enhanced murine cardiomyocyte differentiation via optimized engagement of 3D extracellular matrices, Scientific reports, 5 (2015) 18705.

[8] M.E. Kupfer, W.H. Lin, V. Ravikumar, K. Qiu, L. Wang, L. Gao, D.B. Bhuiyan, M. Lenz, J. Ai, R.R. Mahutga, D. Townsend, J. Zhang, M.C. McAlpine, E.G. Tolkacheva, B.M. Ogle, In Situ Expansion, Differentiation, and Electromechanical Coupling of Human Cardiac Muscle in a 3D Bioprinted, Chambered Organoid, Circ Res, 127 (2020) 207–224.

[9] P.W. Burridge, E. Matsa, P. Shukla, Z.C. Lin, J.M. Churko, A.D. Ebert, F. Lan, S. Diecke, B. Huber, N.M. Mordwinkin, J.R. Plews, O.J. Abilez, B. Cui, J.D. Gold, J.C. Wu, Chemically defined generation of human cardiomyocytes, Nat Methods, 11 (2014) 855–860.

[10] J.C. Boucaut, T. Darribere, Fibronectin in early amphibian embryos. Migrating mesodermal cells contact fibronectin established prior to gastrulation, Cell Tissue Res, 234 (1983) 135–145.

[11] J.C. Boucaut, T. Darribere, H. Boulekbache, J.P. Thiery, Prevention of gastrulation but not neurulation by antibodies to fibronectin in amphibian embryos, Nature, 307 (1984) 364–367.

[12] G. Lee, R. Hynes, M. Kirschner, Temporal and spatial regulation of fibronectin in early Xenopus development, Cell, 36 (1984) 729–740.

[13] K.K. Linask, J.W. Lash, Precardiac cell migration: fibronectin localization at mesoderm-endoderm interface during directional movement, Dev Biol, 114 (1986) 87–101.

[14] T. Darribere, K.M. Yamada, K.E. Johnson, J.C. Boucaut, The 140-kDa fibronectin receptor complex is required for mesodermal cell adhesion during gastrulation in the amphibian Pleurodeles waltlii, Dev Biol, 126 (1988) 182–194.

[15] K.K. Linask, J.W. Lash, A role for fibronectin in the migration of avian precardiac cells. I. Dose-dependent effects of fibronectin antibody, Dev Biol, 129 (1988) 315–323.

[16] K.K. Linask, J.W. Lash, A role for fibronectin in the migration of avian precardiac cells. II. Rotation of the heart-forming region during different stages and its effects, Dev Biol, 129 (1988) 324–329.

[17] K.E. Johnson, T. Darribere, J.C. Boucaut, Mesodermal cell adhesion to fibronectin-rich fibrillar extracellular matrix is required for normal Rana pipiens gastrulation, J Exp Zool, 265 (1993) 40–53.

[18] H.R. Suzuki, M. Solursh, H.S. Baldwin, Relationship between fibronectin expression during gastrulation and heart formation in the rat embryo, Dev Dyn, 204 (1995) 259–277.

[19] J.C. Boucaut, L. Clavilier, T. Darribere, M. Delarue, J.F. Riou, D.L. Shi, What mechanisms drive cell migration and cell interactions in Pleurodeles?, Int J Dev Biol, 40 (1996) 675–683.

[20] J.P. Thiery, J.P. Sleeman, Complex networks orchestrate epithelial-mesenchymal transitions, Nat Rev Mol Cell Biol, 7 (2006) 131–142.

[21] J. Lim, J.P. Thiery, Epithelial-mesenchymal transitions: insights from development, Development, 139 (2012) 3471–3486.

[22] M. Leptin, B. Grunewald, Cell shape changes during gastrulation in Drosophila, Development, 110 (1990) 73–84.

[23] M.A. Nieto, M.G. Sargent, D.G. Wilkinson, J. Cooke, Control of cell behavior during vertebrate development by Slug, a zinc finger gene, Science, 264 (1994) 835–839.

[24] J.P. Thiery, H. Acloque, R.Y. Huang, M.A. Nieto, Epithelial-mesenchymal transitions in development and disease, Cell, 139 (2009) 871–890.

[25] M.A. Nieto, The snail superfamily of zinc-finger transcription factors, Nat Rev Mol Cell Biol, 3 (2002) 155–166.

[26] A. Barrallo-Gimeno, M.A. Nieto, The Snail genes as inducers of cell movement and survival: implications in development and cancer, Development, 132 (2005) 3151–3161.

[27] M. Bachmann, S. Kukkurainen, V.P. Hytonen, B. Wehrle-Haller, Cell Adhesion by Integrins, Physiol Rev, 99 (2019) 1655–1699.

[28] M. Bharadwaj, N. Strohmeyer, G.P. Colo, J. Helenius, N. Beerenwinkel, H.B. Schiller, R. Fassler, D.J. Muller, alphaV-class integrins exert dual roles on alpha5beta1 integrins to strengthen adhesion to fibronectin, Nat Commun, 8 (2017) 14348.

[29] G.E. Hannigan, C. Leung-Hagesteijn, L. Fitz-Gibbon, M.G. Coppolino, G. Radeva, J. Filmus, J.C. Bell, S. Dedhar, Regulation of cell adhesion and anchorage-dependent growth by a new beta 1-integrin-linked protein kinase, Nature, 379 (1996) 91–96.

[30] M. Delcommenne, C. Tan, V. Gray, L. Rue, J. Woodgett, S. Dedhar, Phosphoinositide-3- OH kinase-dependent regulation of glycogen synthase kinase 3 and protein kinase B/AKT by the integrin-linked kinase, Proc Natl Acad Sci U S A, 95 (1998) 11211–11216.

[31] A. Oloumi, T. McPhee, S. Dedhar, Regulation of E-cadherin expression and beta-catenin/Tcf transcriptional activity by the integrin-linked kinase, Biochim Biophys Acta, 1691 (2004) 1–15.

[32] X. Lian, C. Hsiao, G. Wilson, K. Zhu, L.B. Hazeltine, S.M. Azarin, K.K. Raval, J. Zhang, T.J. Kamp, S.P. Palecek, Robust cardiomyocyte differentiation from human pluripotent stem cells via temporal modulation of canonical Wnt signaling, Proc Natl Acad Sci U S A, 109 (2012) E1848–1857.

[33] X. Lian, J. Zhang, S.M. Azarin, K. Zhu, L.B. Hazeltine, X. Bao, C. Hsiao, T.J. Kamp, S.P. Palecek, Directed cardiomyocyte differentiation from human pluripotent stem cells by modulating Wnt/beta-catenin signaling under fully defined conditions, Nat Protoc, 8 (2013) 162–175.

[34] L. Yang, M.H. Soonpaa, E.D. Adler, T.K. Roepke, S.J. Kattman, M. Kennedy, E. Henckaerts, K. Bonham, G.W. Abbott, R.M. Linden, L.J. Field, G.M. Keller, Human cardiovascular progenitor cells develop from a KDR+ embryonic-stem-cell-derived population, Nature, 453 (2008) 524–528.

[35] S.J. Kattman, A.D. Witty, M. Gagliardi, N.C. Dubois, M. Niapour, A. Hotta, J. Ellis, G. Keller, Stage-specific optimization of activin/nodal and BMP signaling promotes cardiac differentiation of mouse and human pluripotent stem cell lines, Cell Stem Cell, 8 (2011) 228–240.

[36] N.C. Dubois, A.M. Craft, P. Sharma, D.A. Elliott, E.G. Stanley, A.G. Elefanty, A. Gramolini, G. Keller, SIRPA is a specific cell-surface marker for isolating cardiomyocytes derived from human pluripotent stem cells, Nat Biotechnol, 29 (2011) 1011–1018.

[37] F.L. Conlon, K.M. Lyons, N. Takaesu, K.S. Barth, A. Kispert, B. Herrmann, E.J. Robertson, A primary requirement for nodal in the formation and maintenance of the primitive streak in the mouse, Development, 120 (1994) 1919–1928.

[38] J. Lough, M. Barron, M. Brogley, Y. Sugi, D.L. Bolender, X. Zhu, Combined BMP-2 and FGF-4, but neither factor alone, induces cardiogenesis in non-precardiac embryonic mesoderm, Dev Biol, 178 (1996) 198–202.

[39] T. Mima, H. Ueno, D.A. Fischman, L.T. Williams, T. Mikawa, Fibroblast growth factor receptor is required for in vivo cardiac myocyte proliferation at early embryonic stages of heart development, Proc Natl Acad Sci U S A, 92 (1995) 467–471.

[40] G. Winnier, M. Blessing, P.A. Labosky, B.L. Hogan, Bone morphogenetic protein-4 is required for mesoderm formation and patterning in the mouse, Genes Dev, 9 (1995) 2105–2116.

[41] M.A. Laflamme, K.Y. Chen, A.V. Naumova, V. Muskheli, J.A. Fugate, S.K. Dupras, H. Reinecke, C. Xu, M. Hassanipour, S. Police, C. O’Sullivan, L. Collins, Y. Chen, E. Minami, E.A. Gill, S. Ueno, C. Yuan, J. Gold, C.E. Murry, Cardiomyocytes derived from human embryonic stem cells in pro-survival factors enhance function of infarcted rat hearts, Nat Biotechnol, 25 (2007) 1015–1024.

[42] J.L. Duband, J.P. Thiery, Appearance and distribution of fibronectin during chick embryo gastrulation and neurulation, Dev Biol, 94 (1982) 337–350.

[43] E.L. George, E.N. Georges-Labouesse, R.S. Patel-King, H. Rayburn, R.O. Hynes, Defects in mesoderm, neural tube and vascular development in mouse embryos lacking fibronectin, Development, 119 (1993) 1079–1091.

[44] E. Spiegel, M. Burger, M. Spiegel, Fibronectin in the developing sea urchin embryo, J Cell Biol, 87 (1980) 309–313.

[45] C. Wu, S.Y. Keightley, C. Leung-Hagesteijn, G. Radeva, M. Coppolino, S. Goicoechea, J.A. McDonald, S. Dedhar, Integrin-linked protein kinase regulates fibronectin matrix assembly, E-cadherin expression, and tumorigenicity, J Biol Chem, 273 (1998) 528–536.

[46] A. Novak, S.C. Hsu, C. Leung-Hagesteijn, G. Radeva, J. Papkoff, R. Montesano, C. Roskelley, R. Grosschedl, S. Dedhar, Cell adhesion and the integrin-linked kinase regulate the LEF-1 and beta-catenin signaling pathways, Proc Natl Acad Sci U S A, 95 (1998) 4374–4379.

[47] A.J. Engler, S. Sen, H.L. Sweeney, D.E. Discher, Matrix elasticity directs stem cell lineage specification, Cell, 126 (2006) 677–689.

[48] T. Rozario, D.W. DeSimone, The extracellular matrix in development and morphogenesis: a dynamic view, Dev Biol, 341 (2010) 126–140.

[49] A. Laperle, C. Hsiao, M. Lampe, J. Mortier, K. Saha, S.P. Palecek, K.S. Masters, alpha-5 Laminin Synthesized by Human Pluripotent Stem Cells Promotes Self-Renewal, Stem Cell Reports, 5 (2015) 195–206.

[50] J. Zhang, G.F. Wilson, A.G. Soerens, C.H. Koonce, J. Yu, S.P. Palecek, J.A. Thomson, T.J. Kamp, Functional cardiomyocytes derived from human induced pluripotent stem cells, Circ Res, 104 (2009) e30–41.

